# Optimising a coordinate ascent algorithm for the meta-analysis of test accuracy studies

**DOI:** 10.1101/2022.12.05.519131

**Authors:** Mohammed Baragilly, Brian H Willis

## Abstract

Meta-analysis may be used to summarise a test’s accuracy. Often the sensitivity and specificity are the measures of interest and as these are correlated a bivariate random effects model is commonly used to fit the data. This model has five parameters and it may be optimised using a Newton-Raphson based algorithm providing adequate initial values of the parameters are identified. Numerical methods may be used to estimate robust initial values but estimating these is computationally expensive and it is not clear whether they provide a significant advantage over closed form methods in terms of reducing bias, mean square error, average relative error, and coverage probability. Here we consider six closed form methods for estimating the initial values of the parameters for a co-ordinate ascent algorithm used to fit the bivariate model and compare them with numerically derived robust initial values. Using simulation studies we demonstrate that all the closed form methods lead to a reduction in computation time of around 80% and rank higher overall across the metrics when compared with the robust initial values method. Although no initial values estimator dominated the others across all parameters and metrics, the two-step Hedges-Olkin estimator ranked highest overall across the different scenarios.

## 1. Introduction

Test accuracy studies are important in the evaluation of diagnostic technologies [1,2] and when there are multiple primary studies evaluating the same test these may be aggregated using meta-analysis [3, 4]. The sensitivity and specificity are the two main statistics used to summarise a test’s performance and these depend on the threshold for a positive test result. Even when the threshold is constant between studies, they are still likely to be correlated and this is assumed in many of the models used in meta-analysis [3,4].

The bivariate random effects model (BRM) is one of most common models used to pool test accuracy studies when the data are reported at a constant threshold [4, Chu and Cole 2006]. There are five parameters to estimate and as the likelihood function has no closed form these must be estimated using numerical methods.

A few functions and packages exist in R [5] SAS [6], and Stata [7] which may be used to fit the BRM. However, as they are designed to fit a range of generalised linear mixed models and not specifically optimised for the BRM this may affect probability of convergence when fitting the BRM. For example, a simulation study demonstrated the *glmer* function from the lme4 package in R had a probability of convergence of less than 85% when fitting the BRM [8].

For many optimisation methods the initial values of the parameters can be important in determining convergence on a global maximum [9]. However, specifically for Newton-Raphson methods, Kantorovich’s theorem asserts that the Newton-Raphson algorithm will converge on the root to a function near some initial point providing the Jacobian of the function at the initial point is Lipschitz continuous and its inverse satisfies certain boundedness conditions [10].

Here we evaluate how different methods for estimating the initial values may affect the performance of a Newton-Raphson based co-ordinate ascent algorithm for the BRM. Broadly such methods divide into numerical and analytical methods. For the former we consider robust initial values, which are estimated after deriving a feasible hyperspace for the unknown parameters. As this involves an iterative process, their estimation can be computationally intensive. Furthermore robust initial values do not always lead to unbiased results in the final estimate for the parameters [10].

In contrast to numerical methods, analytical methods have the advantage of having explicit formulae so do not require estimation by numerical iterative methods thereby increasing computational efficiency. Potentially analytical approaches such as those based on the method of moments [11] may be used to derive initial estimates for the parameters for input to the optimisation algorithm. It is also not clear how these methods compare with numerical methods in other characteristics such as convergence, bias, mean squared error and coverage.

Here six approaches based on the method of moments will be explored [12-16] for their potential to provide initial values for the co-ordinate ascent algorithm for the likelihood of the BRM. Using a simulation study these are compared with robust initial values in terms of the mean bias, mean squared error, coverage probability and mean computation time.

The paper is organised as follows. In section 2, we discuss the bivariate random effects model used in test accuracy meta-analyses. In section 3, a co-ordinate ascent algorithm for the likelihood based on the Newton-Raphson method is discussed. In sections 4 and 5, we describe numerical and analytical methods that may be used to derive initial starting values for the algorithm used to fit the BRM. In sections 6 and 7, these methods are compared using a simulation study and applying them to two real datasets from the literature. In section 8, we end with the discussion and conclusion.

## 2. The bivariate model

The BRM jointly analyses the sensitivity and specificity by assuming independent binomial distributions for the true positives, and true negatives within each study and a bivariate normal distribution for the logits of the sensitivity and specificity between studies [4]. Thus we have a model of the form

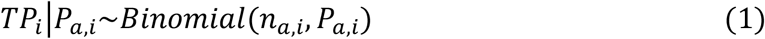

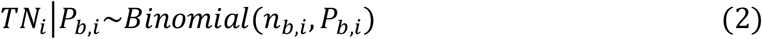

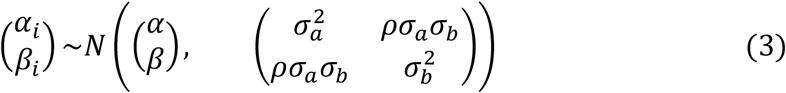

where for the i^th^ study, *TP*_*i*_, *TN*_*i*_, *n*_*a,i*_ and *n*_*b,i*_ are the number of true positives, true negatives, diseased, and non-diseased. The study-level parameters for the sensitivity and specificity are *P*_*a,i*_ and *P*_*b,i*_ respectively and *α*_*i*_ = *logit*(*P*_*a,i*_) and *β*_*i*_ = *logit*(*P*_*b,i*_). For the bivariate normal distribution, *α* and 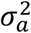 are the mean and variance for the logit sensitivities, *β* and 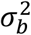 are the mean and variance for the logit specificities, and *ρ* is the correlation between *α*_*i*_ and *β*_*i*_ across studies, respectively.

In order to make inferences on both the sensitivity and specificity, the five parameters 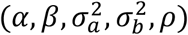 of the bivariate generalized linear mixed effect model need to be estimated in advance. The maximum likelihood estimates (MLE) can be derived by maximising the log likelihood function

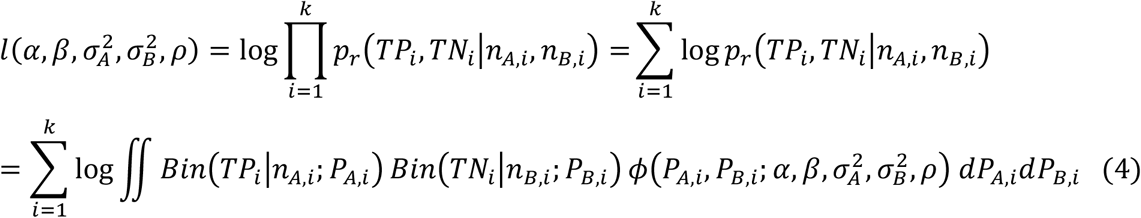

where

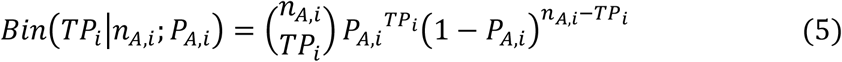

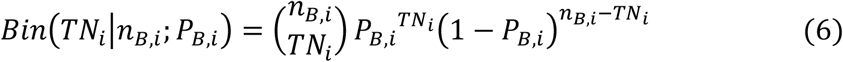

and 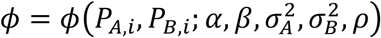 is the bivariate logit normal distribution, such that

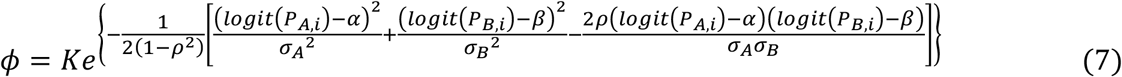

where

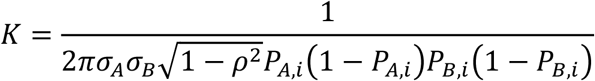

## 3. A coordinate ascent algorithm for the BRM

The likelihood function for the BRM in (4) does not have a closed form so the challenge is to find the maximum likelihood estimates numerically for the 5 parameters 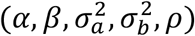. Although these parameters may be estimated simultaneously using algorithms based on the Fisher Information for example, the method of profiling partitions the parameters into those of interest and nuisance parameters and maximises over the latter. This profile likelihood function of the parameters of interest is then maximised to give the MLE for the parameter(s) of interest. Similarly the MLEs for the other parameters may be obtained.

The coordinate ascent algorithm further simplifies estimating the 5 parameters of the BRM. Essentially this focusses on one parameter at a time and holds the remaining parameters constant when maximising the log likelihood with respect to the one parameter permitted to vary. This is done for each of the parameters, the parameter estimates are iteratively updated and the algorithm keeps cycling through the parameters until convergence is achieved.

Maximising the log-likelihood, 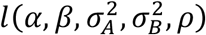, entails finding the root of its derivative with respect to the parameter of interest. If 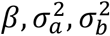 and *ρ* are held fixed and *α* is allowed to vary, an updated estimate 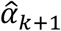 for *α* may be obtained as follows. Let 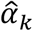 be the last estimate for *α* and set 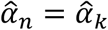 then iterate 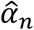 using the Newton-Raphson method as follows

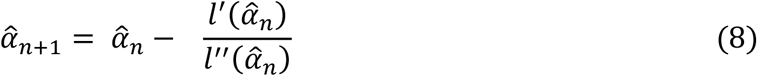

where the first and second derivatives may be estimated numerically from

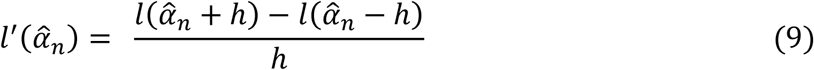

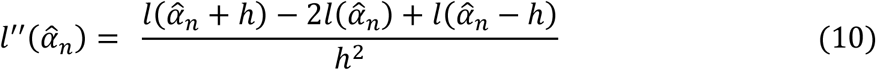

The log-likelihood function in (4) contains a double integral and this is estimated for each iteration. The integral may be estimated numerically using adaptive Gaussian quadrature which may be found in the *adaptIntegrate* function in R [17]. The iterations are continued until convergence is achieved, when the condition 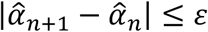 is satisfied for *ε* = 10^−6^. The updated estimate 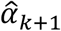 is set to the converged estimate, 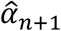 and this remains the fixed estimate for *α* whilst the process is repeated for the other parameters.

## 4. Estimating initial values for the coordinate ascent algorithm

In the algorithm discussed in section 3, the Newton-Raphson method is used to estimate the parameters in the coordinate ascent algorithm. Although, in general, initial values are important to iterative algorithms, Kantorovich’s theorem reinforces the importance of the choice of initial values for a Newton-Raphson algorithm if it is to converge [10]. There are a number of methods that potentially could be used to estimate the initial values for the algorithm including numerical and analytical approaches.

A numerical approach that can be used involves defining a feasible space for the global maximum of the parameter vector, 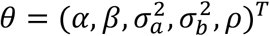. The feasible space is based on the minimum and maximum values for the sensitivity, specificity, and associated variances across the studies. An equally spaced grid is overlaid on the feasible space [18] and *ρ* is set to zero. Thus, all the points on the grid in five-dimensional hyperspace represent candidates for the initial values for the parameter vector, 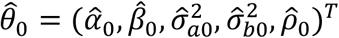. This approach provides maximum likelihood robust initial start values (MLRIV) and has been used previously [8].

A major drawback with estimating robust initial values is that it is computationally intensive. Furthermore it is not clear whether it offers a significant advantage over other more direct methods for estimating the initial values. These more direct approaches are described in the next section.

## 5. Analytical estimation of the initial starts in the coordinate ascent algorithm

For a univariate meta-analysis of *k* studies the unknown parameter is usually estimated by

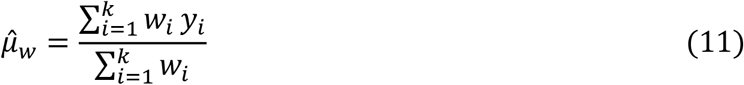

where *y*_*i*_ is the observed effect for the i^th^ study and *w*_*i*_ is the associated weight for the study [19]. Usually the weights are derived from the inverse variance as this maximises the likelihood function and under the assumptions of a random effects model the variance includes both a within-study and between-study component. This gives *w*_*i,RE*_ = *1/*(*V*_*i*_ + *τ*^2^) where *V*_*i*_ is the within-study variance and *τ*^2^ is the between-study variance [19].

For the bivariate random effects model we may use estimates derived from two univariate meta-analyses to provide the initial values for the coordinate ascent algorithm. Thus to estimate 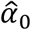, for sensitivity, *Sen*, we have *y*_*i*_ = *logit*(*Sen*_*i*_) and the within-study variance *V*_*i*_ = *1/n*_*i*_*Sen*_*i*_(*1 − Sen*_*i*_) estimated by the delta method, where *n*_*i*_ is the number of diseased in study *i* [4].

However, central to estimating 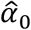 is being able to derive an estimate for the associated between-study variance 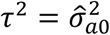. Similarly for 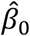. In the next six sub-sections all the methods described are closed form methods used to estimate the between-study variance *τ*^2^.

It also should be assumed in the methods that follow that for the *i*^*th*^ study, *y*_*i*_ refers to either the *logit*(*Sensitivity*_*i*_) or *logit*(*Specificity*_*i*_) and *V*_*i*_ is the corresponding variance estimated by the delta method [4]. An extensive review on the methods that follow may be found in Veroniki et al [20].

### 5.1 DerSimonian and Laird (DL) method

In this approach, the estimate for *τ*^2^ is derived using a technique based on the method of moments, by equating the Q statistic to its expected value and solving for *τ*^2^ [12]. Generally, for individual study weights, *a*_*i*_ and associated variances, *V*_*i*_ the general form for the estimator, 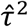 is given by

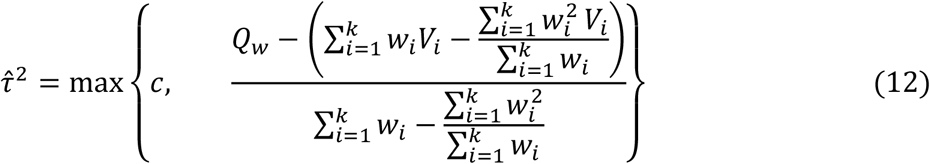

where 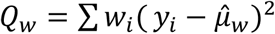 [19]. When the weights, *w*_*i*_ are set to the *1/V*_*i*_, the inverse variance, the term in brackets in the numerator reduces to *k* – 1 and this gives the DerSimonian and Laird estimator, 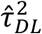[12]. In general *c* is usually set to zero.

### 5.2 Two-step estimator with DerSimonian and Laird (DL2) method

The estimates for 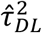 and *Q*_*w*_ are derived by assuming an initial fixed effects model, so that the weights *w*_*i*_= *1/V*_*i*_, where *V*_*i*_ are the individual study variances.

Once an estimate for the between study variance, *τ*^2^ has been obtained the individual study weights may be modified to include the between-study variance to give the weights of a random effects model, 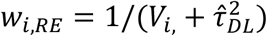. These may be used to give the two-step estimator of DerSimonian and Laird, 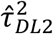 [13].

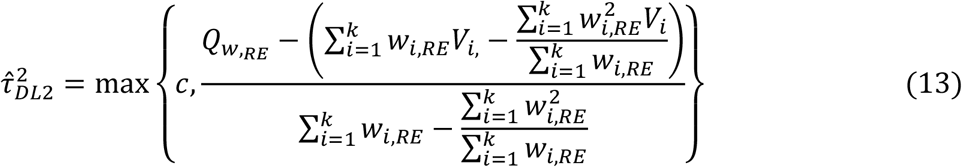

where 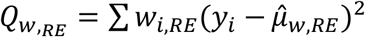.

### 5.3 Hedges and Olkin (HO) method

The Hedges and Olkin estimator [14] was in fact first proposed by Cochran [20]. Although it is derived using the methods of moments, it may be easily deduced from (12) by setting *w*_*i*_ = *1/k* [13, 20] giving

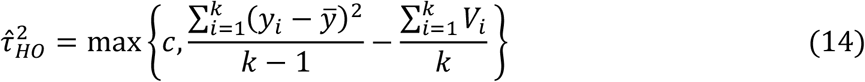

### 5.4 Two-step estimator for the Hedges and Olkin (HO2) method

Like the DL2 method [13], the two-step Hedges and Olkin estimator HO2 requires first to get the estimator from the fixed effects model, and then use it to get the random effects weights as a second step. Thus, using weights, 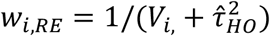, we re-estimate 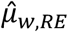 and 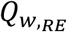 and these are substituted into equation (13) to give 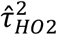 [13].

Previous analyses have shown that, in general, the two-step estimators HO2 and DL2 outperform the one-step HO and DL estimators, and that the two-step estimators are preferred over their one-step counterparts when non-iterative methods are used to estimate the between-study variance [13].

### 5.5 Hartung and Makambi (HM) method

Without imposing a lower limit of *c = 0*, the DL and HO estimators potentially may return negatives values. The main idea behind HM estimator [15, 20] is to use the quadratic form of the random variables and multiply it by the factor *Q/*(*2*(*k* − *1)* + *Q)* to ensure positivity rather than truncating the values to zero.

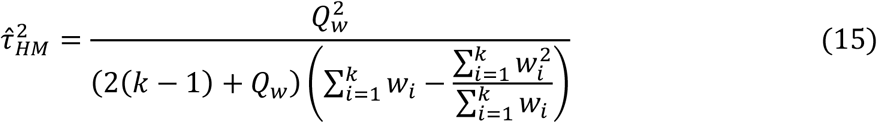

where 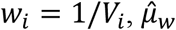 is as estimated in (11) and 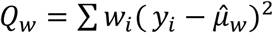.

### 5.6 Hunter and Schmidt (HS) method

The HS estimator as introduced by Hunter and Schmidt in 2004 [16] was proposed to measure the level of heterogeneity when analysing correlation coefficients [16]. This has been demonstrated to be easily generalised to other effect measures [22] and can be obtained from,

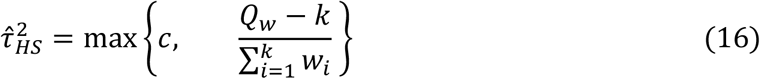

where *w*_*i*_ = *1/V*_*i*_ and *Q*_*w*_ is defined as previously.

From a simulation study, Kontopantelis et al [23] concluded that setting *c* = 0.01 resulted in lower bias for truncated estimators of between-study variance than when c = 0. From now on, for the simulation study that follows *c* is set to 0.01 for all of the truncated estimators.

### 5.7 Correlation between the between-study variances

In order to get the estimated initial value of the correlation term between *α* = *logit*(*Sensitivity)* and *β* = *logit*(*Specificity)* we can estimate the Pearson correlation coefficient from the observed data. Let *lse*_*i*_ and *lsp*_*i*_ be the logits of the observed sensitivity and specificity for the *i*^*th*^ study, and *m*_*lSe*_ and *m*_*lsp*_ be their unweighted means respectively. Then an initial estimate for the correlation, 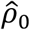 is given by

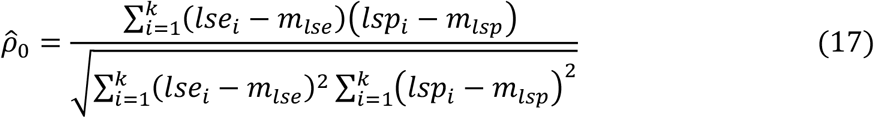

Accordingly, we may use the random effects weights, *w*_*i,RE*_ to estimate 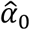 and 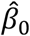; each of (12) – (16) to give different estimates for 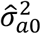 and 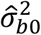; and (17) to estimate 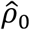. These will provide the initial estimates for the parameter vector 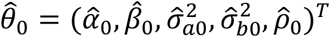 that will be used in the optimisation algorithm.

In the next section, the effects the different estimators for the initial values have on the performance of the co-ordinate ascent algorithm for fitting the BRM are compared with robust initial values.

## 6. Simulation study

Here, the co-ordinate ascent algorithm based on the six initial estimates of 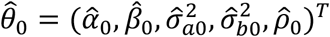 is evaluated through a simulated study. They are compared with the algorithm based on the MLRIV [8]. The section ends with two real case studies. The analysis here is conducted in R [24].

For the simulations studies, the datasets were generated by simulating from the BRM in (4) with the condition that the true values of the logit sensitivity *α* and logit specificity *β* were randomly simulated to satisfy the *f < s* and varying the: (i) Numbers of studies, *k* = {5, 10, 15, 20, 25 }for 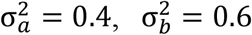, and *ρ* = −0.7; (ii) Correlation, *ρ* ={-0.1, -0.3, -0.5, - 0.9} for 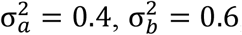, and *k*=10; (iii) Variances, 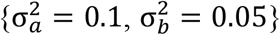 and 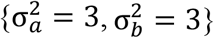 for *ρ* = −0.5 and *k*=10.

This provides the study-specific sensitivity, *P*_*A,i*_ = *logit*^−1^(*α*_i_) and specificity *P*_*B,i*_ = *logit*^−1^(*β*_i_). For each study *i*, the number of non-diseased *n*_*b,i*_, was generated randomly to be between 50 and 1000 and the diseased *n*_*a,i*_, chosen to be *γn*_*b,i*_ rounded to the nearest whole number, where γ was randomly simulated to be between 0.05 and 0.5. Thus, for each of *k* studies, the true positives *TP*_*i*_, and true negatives *TN*_*i*_ were simulated from the binomial distributions detailed in (1) and (2).

For each of the initial estimators, the coordinate ascent algorithm to the BRM was applied to 10,000 simulated data sets. The results were compared using the following performance metrics: the mean squared error (MSE), mean bias, average relative error (ARE), coverage probability and average computing time in minutes. For the average relative error (ARE), we used the formula:

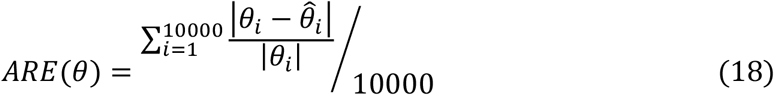

where 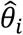 is the estimated value of the true value *θ*_*i*_ in the *i*-th iteration.

The estimators were ranked 1 to 7 for each of the MSE, absolute bias and ARE over each of the parameters. In order to give an overall comparison, the rank scores were aggregated to give an overall score for each estimator. These were then ranked from lowest to highest.

Tables 1 to 5 give the performance metrics for each of the initial estimators for *k=*5, 10, 15, 20 and 25 studies respectively. Tables 6 to 9 give the performance metrics for each of the initial estimators for *ρ* = -0.1, -0.3, -0.5, -0.9. Tables 10 and 11 give the performance metrics for each of the initial estimators for a ‘homogenous’ population, (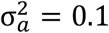 and 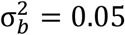) and a ‘heterogeneous’ population, (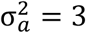 and 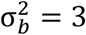). The distributions for each of the parameters are given as box and whisker plots in the appendix.

**Table 1:** MSE, mean bias, ARE, average computing time in minutes and average total iteration of the estimated values of the five parameters using coordinate ascent algorithm based on different initial starts at *k*=5.

| Criteria | DL | DL2 | HO | HO2 | HM | HS | MLRIV |
| --- | --- | --- | --- | --- | --- | --- | --- |
| $MSE(\hat{\alpha})$ | 2.6238 | 2.6807 | 2.6782 | 2.7986 | 2.7028 | 2.6996 | 2.9015 |
| $MSE(\hat{\beta})$ | 2.5144 | 2.5174 | 2.5321 | 2.5495 | 2.5312 | 2.5456 | 2.6381 |
| $MSE(\hat{\sigma}_a^2)$ | 0.2255 | 0.2622 | 0.3386 | 0.2735 | 0.2132 | 0.2358 | 1.3274 |
| $MSE(\hat{\sigma}_b^2)$ | 0.3173 | 0.2893 | 0.3086 | 0.2564 | 0.2469 | 0.2783 | 0.6045 |
| $MSE(\hat{\rho})$ | 0.1560 | 0.1523 | 0.1500 | 0.1513 | 0.1756 | 0.1466 | 0.2362 |
| $bias(\hat{\alpha})$ | -0.0011 | -0.0060 | 0.0011 | 0.0030 | 0.0038 | 0.0035 | 0.0280 |
| $bias(\hat{\beta})$ | 0.0057 | 0.0019 | -0.0031 | 0.0042 | 0.0003 | 0.0003 | 0.0167 |
| $bias(\hat{\sigma}_a^2)$ | -0.0708 | -0.0544 | -0.0711 | -0.0690 | -0.0558 | -0.0802 | 0.1679 |
| $bias(\hat{\sigma}_b^2)$ | -0.0978 | -0.1019 | -0.0986 | -0.1046 | -0.1133 | -0.1247 | -0.0773 |
| $bias(\hat{\rho})$ | 0.0959 | 0.0889 | 0.0891 | 0.0913 | 0.1141 | 0.0864 | 0.1843 |
| $ARE(\hat{\alpha})$ | 1.8642 | 76.3056 | 1.7070 | 1.5432 | 2.6680 | 1.4761 | 1.6085 |
| $ARE(\hat{\beta})$ | 1.2666 | 0.8701 | 1.2561 | 0.9510 | 2.2908 | 0.9517 | 0.9678 |
| $ARE(\hat{\sigma}_a^2)$ | 0.7898 | 0.8150 | 0.8163 | 0.8000 | 0.7727 | 0.7975 | 1.0492 |
| $ARE(\hat{\sigma}_b^2)$ | 0.6192 | 0.6071 | 0.6152 | 0.6050 | 0.6072 | 0.6211 | 0.6573 |
| $ARE(\hat{\rho})$ | 0.3838 | 0.3793 | 0.3788 | 0.3781 | 0.4047 | 0.3726 | 0.4588 |
| Total rank score | 56 | 52 | 57 | 53 | 59 | 49 | 92 |
| Overall rank | 4 | 2 | 5 | 3 | 6 | 1 | 7 |
| Coverage probability | 0.9825 | 0.9820 | 0.9813 | 0.9808 | 0.9815 | 0.9796 | 0.9774 |
| Av. computing time | 8.5225 | 8.6408 | 8.4370 | 7.9489 | 9.6272 | 10.8305 | 33.691 |

**Table 2:** MSE, mean bias, ARE, average computing time in minutes and coverage probability of the estimated values of the five parameters using co-ordinate ascent algorithm based on different initial starts at *k*=10.

| Criteria | DL | DL2 | HO | HO2 | HM | HS | MLRIV |
| --- | --- | --- | --- | --- | --- | --- | --- |
| $MSE(\hat{\alpha})$ | 2.6674 | 2.4995 | 2.5299 | 2.6111 | 2.6065 | 2.6558 | 2.9148 |
| $MSE(\hat{\beta})$ | 2.4682 | 2.4261 | 2.3701 | 2.4234 | 2.4192 | 2.4388 | 2.9387 |
| $MSE(\hat{\sigma}_a^2)$ | 0.1239 | 0.1232 | 0.1294 | 0.1521 | 0.1722 | 0.1435 | 0.9826 |
| $MSE(\hat{\sigma}_b^2)$ | 0.1434 | 0.1580 | 0.1596 | 0.1460 | 0.1500 | 0.1473 | 0.4522 |
| $MSE(\hat{\rho})$ | 0.0680 | 0.0696 | 0.0680 | 0.0718 | 0.0877 | 0.0677 | 0.1444 |
| $bias(\hat{\alpha})$ | -0.0014 | 0.0031 | 0.0070 | 0.0031 | -0.0031 | 0.0014 | 0.0232 |
| $bias(\hat{\beta})$ | -0.0052 | -0.0036 | -0.0068 | -0.0014 | 0.0032 | -0.0013 | 0.0195 |
| $bias(\hat{\sigma}_a^2)$ | -0.0536 | -0.0495 | -0.0537 | -0.0501 | -0.0391 | -0.0452 | 0.1389 |
| $bias(\hat{\sigma}_b^2)$ | -0.0706 | -0.0646 | -0.0681 | -0.0712 | -0.0724 | -0.0759 | -0.0294 |
| $bias(\hat{\rho})$ | 0.0494 | 0.0483 | 0.0472 | 0.0504 | 0.0672 | 0.0468 | 0.1308 |
| $ARE(\hat{\alpha})$ | 1.0251 | 1.0964 | 1.0393 | 1.4473 | 1.1749 | 2.6287 | 2.1397 |
| $ARE(\hat{\beta})$ | 0.6148 | 0.7682 | 0.8410 | 0.7888 | 0.6665 | 0.9773 | 0.7350 |
| $ARE(\hat{\sigma}_a^2)$ | 0.6390 | 0.6355 | 0.6309 | 0.6483 | 0.6301 | 0.6479 | 0.8574 |
| $ARE(\hat{\sigma}_b^2)$ | 0.4636 | 0.4634 | 0.4620 | 0.4688 | 0.4592 | 0.4662 | 0.5004 |
| $ARE(\hat{\rho})$ | 0.2611 | 0.2631 | 0.2597 | 0.2666 | 0.2857 | 0.2599 | 0.3494 |
| Total rank score | 49 | 47 | 50 | 65 | 54 | 56 | 94 |
| Overall rank | 2 | 1 | 3 | 6 | 4 | 5 | 7 |
| Coverage probability | 0.9455 | 0.9457 | 0.9456 | 0.9466 | 0.9442 | 0.9482 | 0.9482 |
| Av. computing time | 11.5817 | 12.4905 | 11.8547 | 12.7100 | 12.2705 | 13.4736 | 65.9456 |

**Table 3:** MSE, mean bias, ARE, average computing time in minutes and coverage probability of the estimated values of the five parameters using co-ordinate ascent algorithm based on different initial starts at *k*=15.

| Criteria | DL | DL2 | HO | HO2 | HM | HS | MLRIV |
| --- | --- | --- | --- | --- | --- | --- | --- |
| $MSE(\hat{\alpha})$ | 2.5048 | 2.5874 | 2.5004 | 2.5388 | 2.6122 | 2.5793 | 2.7651 |
| $MSE(\hat{\beta})$ | 2.4188 | 2.5212 | 2.4699 | 2.4106 | 2.3900 | 2.4691 | 2.5892 |
| $MSE(\hat{\sigma}_a^2)$ | 0.1310 | 0.1219 | 0.1066 | 0.1213 | 0.1114 | 0.0965 | 1.0535 |
| $MSE(\hat{\sigma}_b^2)$ | 0.1312 | 0.1448 | 0.1315 | 0.1332 | 0.1260 | 0.1245 | 0.3249 |
| $MSE(\hat{\rho})$ | 0.0482 | 0.0452 | 0.0467 | 0.0471 | 0.0555 | 0.0471 | 0.0900 |
| $bias(\hat{\alpha})$ | -0.0011 | -0.0005 | -0.0030 | -0.0019 | -0.0019 | 0.0024 | 0.0197 |
| $bias(\hat{\beta})$ | -0.0026 | 0.0004 | -0.0028 | -0.0007 | 0.0032 | -0.0025 | 0.0124 |
| $bias(\hat{\sigma}_a^2)$ | -0.0280 | -0.0331 | -0.0322 | -0.0334 | -0.0286 | -0.0378 | 0.1264 |
| $bias(\hat{\sigma}_b^2)$ | -0.0449 | -0.0379 | -0.0430 | -0.0481 | -0.0434 | -0.0483 | -0.0055 |
| $bias(\hat{\rho})$ | 0.0294 | 0.0289 | 0.0314 | 0.0305 | 0.0409 | 0.0301 | 0.0899 |
| $ARE(\hat{\alpha})$ | 1.0634 | 4.9751 | 1.2252 | 1.0373 | 0.8177 | 0.9212 | 0.9591 |
| $ARE(\hat{\beta})$ | 0.5432 | 1.1189 | 0.6492 | 0.7901 | 1.3111 | 0.5958 | 1.0349 |
| $ARE(\hat{\sigma}_a^2)$ | 0.5718 | 0.5652 | 0.5583 | 0.5677 | 0.5475 | 0.5523 | 0.7725 |
| $ARE(\hat{\sigma}_b^2)$ | 0.3906 | 0.3990 | 0.3924 | 0.3887 | 0.3957 | 0.3939 | 0.4269 |
| $ARE(\hat{\rho})$ | 0.2244 | 0.2202 | 0.2215 | 0.2235 | 0.2348 | 0.2224 | 0.2799 |
| Total rank score | 52 | 56 | 53 | 55 | 59 | 50 | 93 |
| Overall rank | 2 | 5 | 3 | 4 | 6 | 1 | 7 |
| Coverage probability | 0.9405 | 0.9389 | 0.9402 | 0.9392 | 0.9437 | 0.9363 | 0.9442 |
| Av. computing time | 16.7206 | 17.2128 | 15.7490 | 14.5151 | 15.1068 | 20.0244 | 96.1566 |

**Table 4:**
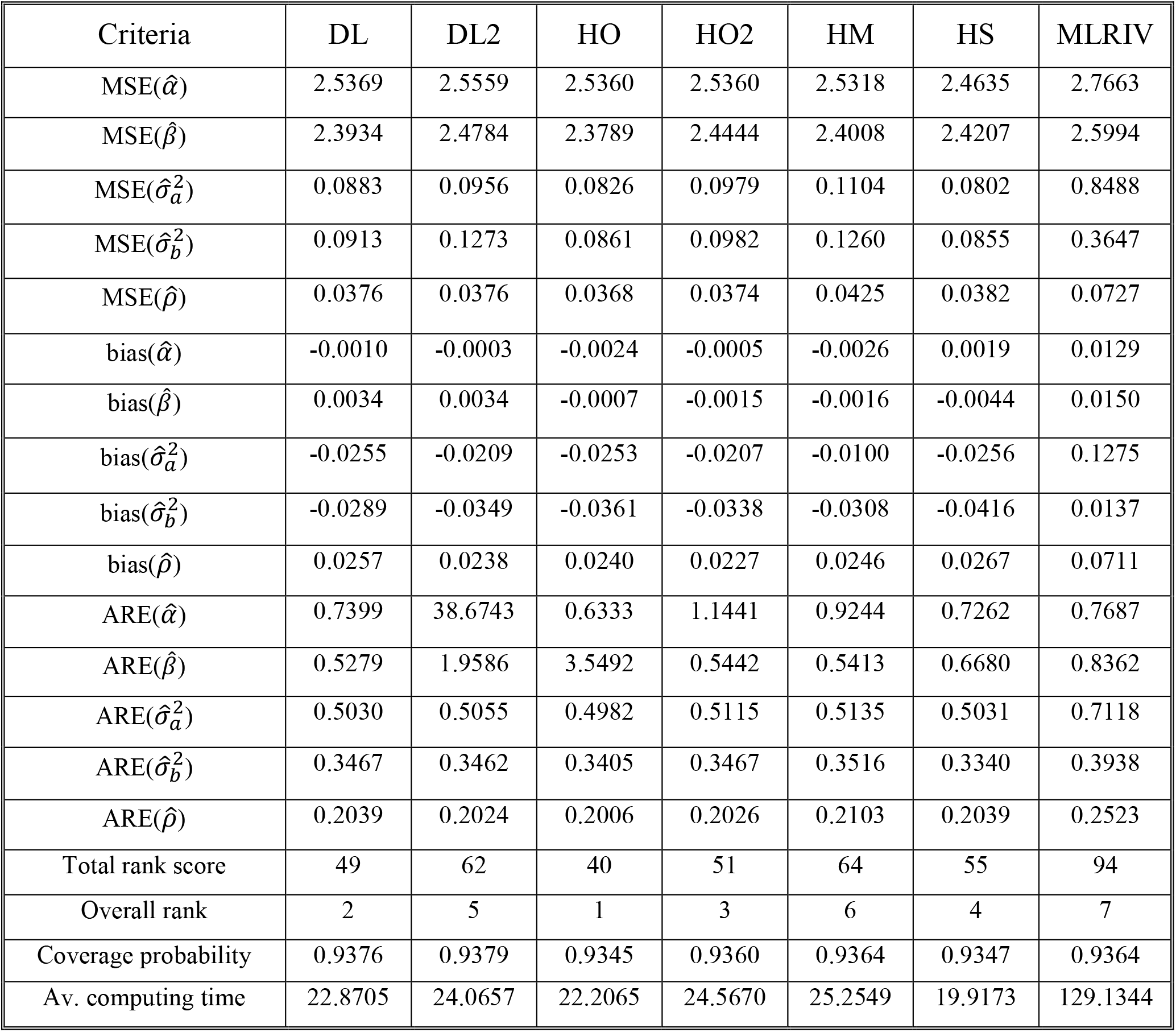
MSE, mean bias, ARE, average computing time in minutes and coverage probability of the estimated values of the five parameters using co-ordinate ascent algorithm based on different initial starts at *k*=20.

| Criteria | DL | DL2 | HO | HO2 | HM | HS | MLRIV |
| --- | --- | --- | --- | --- | --- | --- | --- |
| $MSE(\hat{\alpha})$ | 2.5369 | 2.5559 | 2.5360 | 2.5360 | 2.5318 | 2.4635 | 2.7663 |
| $MSE(\hat{\beta})$ | 2.3934 | 2.4784 | 2.3789 | 2.4444 | 2.4008 | 2.4207 | 2.5994 |
| $MSE(\hat{\sigma}_a^2)$ | 0.0883 | 0.0956 | 0.0826 | 0.0979 | 0.1104 | 0.0802 | 0.8488 |
| $MSE(\hat{\sigma}_b^2)$ | 0.0913 | 0.1273 | 0.0861 | 0.0982 | 0.1260 | 0.0855 | 0.3647 |
| $MSE(\hat{\rho})$ | 0.0376 | 0.0376 | 0.0368 | 0.0374 | 0.0425 | 0.0382 | 0.0727 |
| $bias(\hat{\alpha})$ | -0.0010 | -0.0003 | -0.0024 | -0.0005 | -0.0026 | 0.0019 | 0.0129 |
| $bias(\hat{\beta})$ | 0.0034 | 0.0034 | -0.0007 | -0.0015 | -0.0016 | -0.0044 | 0.0150 |
| $bias(\hat{\sigma}_a^2)$ | -0.0255 | -0.0209 | -0.0253 | -0.0207 | -0.0100 | -0.0256 | 0.1275 |
| $bias(\hat{\sigma}_b^2)$ | -0.0289 | -0.0349 | -0.0361 | -0.0338 | -0.0308 | -0.0416 | 0.0137 |
| $bias(\hat{\rho})$ | 0.0257 | 0.0238 | 0.0240 | 0.0227 | 0.0246 | 0.0267 | 0.0711 |
| $ARE(\hat{\alpha})$ | 0.7399 | 38.6743 | 0.6333 | 1.1441 | 0.9244 | 0.7262 | 0.7687 |
| $ARE(\hat{\beta})$ | 0.5279 | 1.9586 | 3.5492 | 0.5442 | 0.5413 | 0.6680 | 0.8362 |
| $ARE(\hat{\sigma}_a^2)$ | 0.5030 | 0.5055 | 0.4982 | 0.5115 | 0.5135 | 0.5031 | 0.7118 |
| $ARE(\hat{\sigma}_b^2)$ | 0.3467 | 0.3462 | 0.3405 | 0.3467 | 0.3516 | 0.3340 | 0.3938 |
| $ARE(\hat{\rho})$ | 0.2039 | 0.2024 | 0.2006 | 0.2026 | 0.2103 | 0.2039 | 0.2523 |
| Total rank score | 49 | 62 | 40 | 51 | 64 | 55 | 94 |
| Overall rank | 2 | 5 | 1 | 3 | 6 | 4 | 7 |
| Coverage probability | 0.9376 | 0.9379 | 0.9345 | 0.9360 | 0.9364 | 0.9347 | 0.9364 |
| Av. computing time | 22.8705 | 24.0657 | 22.2065 | 24.5670 | 25.2549 | 19.9173 | 129.1344 |

**Table 5:** MSE, mean bias, ARE, average computing time in minutes and coverage probability of the estimated values of the five parameters using co-ordinate ascent algorithm based on different initial starts at *k*=25.

| Criteria | DL | DL2 | HO | HO2 | HM | HS | MLRIV |
| --- | --- | --- | --- | --- | --- | --- | --- |
| $MSE(\hat{\alpha})$ | 2.5517 | 2.5827 | 2.5144 | 2.4528 | 2.4504 | 2.6220 | 2.6329 |
| $MSE(\hat{\beta})$ | 2.4700 | 2.4449 | 2.4105 | 2.4046 | 2.3516 | 2.4775 | 2.5433 |
| $MSE(\hat{\sigma}_a^2)$ | 0.0794 | 0.0890 | 0.0975 | 0.0893 | 0.0945 | 0.0897 | 0.4689 |
| $MSE(\hat{\sigma}_b^2)$ | 0.0937 | 0.0965 | 0.1225 | 0.1106 | 0.0966 | 0.0937 | 0.1851 |
| $MSE(\hat{\rho})$ | 0.0307 | 0.0325 | 0.0306 | 0.0312 | 0.0374 | 0.0308 | 0.0586 |
| $bias(\hat{\alpha})$ | 0.0034 | -0.0005 | -0.0009 | -0.0005 | -0.0012 | -0.0001 | 0.0133 |
| $bias(\hat{\beta})$ | -0.0043 | -0.0027 | -0.0045 | -0.0004 | 0.0062 | -0.0023 | 0.0103 |
| $bias(\hat{\sigma}_a^2)$ | -0.0182 | -0.0196 | -0.0186 | -0.0156 | -0.0091 | -0.0163 | 0.1014 |
| $bias(\hat{\sigma}_b^2)$ | -0.0224 | -0.0246 | -0.0224 | -0.0232 | -0.0212 | -0.0255 | 0.0037 |
| $bias(\hat{\rho})$ | 0.0131 | 0.0161 | 0.0128 | 0.0153 | 0.0201 | 0.0132 | 0.0604 |
| $ARE(\hat{\alpha})$ | 1.8917 | 0.9108 | 1.1304 | 0.6209 | 0.6160 | 0.8001 | 6.2500 |
| $ARE(\hat{\beta})$ | 0.8396 | 0.7995 | 0.5064 | 2.4757 | 0.3754 | 0.5450 | 0.5185 |
| $ARE(\hat{\sigma}_a^2)$ | 0.4661 | 0.4653 | 0.4720 | 0.4702 | 0.4714 | 0.4714 | 0.6246 |
| $ARE(\hat{\sigma}_b^2)$ | 0.3152 | 0.3125 | 0.3202 | 0.3204 | 0.3175 | 0.3116 | 0.3339 |
| $ARE(\hat{\rho})$ | 0.1868 | 0.1887 | 0.1858 | 0.1873 | 0.1982 | 0.1871 | 0.2328 |
| Total rank score | 51 | 58 | 56 | 52 | 53 | 51 | 95 |
| Overall rank | 1 | 6 | 5 | 3 | 4 | 1 | 7 |
| Coverage probability | 0.9315 | 0.9285 | 0.9353 | 0.9344 | 0.9327 | 0.9303 | 0.9381 |
| Av. computing time | 26.0478 | 27.2007 | 27.1192 | 31.6333 | 25.4394 | 31.1322 | 147.1156 |

**Table 6:** MSE, mean bias, ARE, average computing time in minutes and average total iteration of the estimated values of the five parameters using the coordinate ascent algorithm based on different initial starts at *ρ*=-0.1.

| Criteria | DL | DL2 | HO | HO2 | HM | HS | MLRIV |
| --- | --- | --- | --- | --- | --- | --- | --- |
| $MSE(\hat{\alpha})$ | 2.5419 | 2.5194 | 2.5144 | 2.5538 | 2.5047 | 2.5057 | 2.8265 |
| $MSE(\hat{\beta})$ | 2.4867 | 2.4345 | 2.4768 | 2.3250 | 2.4792 | 2.4625 | 2.5701 |
| $MSE(\hat{\sigma}_a^2)$ | 0.1248 | 0.1148 | 0.1185 | 0.1192 | 0.1037 | 0.0968 | 1.1275 |
| $MSE(\hat{\sigma}_b^2)$ | 0.1399 | 0.1318 | 0.1228 | 0.1250 | 0.1271 | 0.1203 | 0.4338 |
| $MSE(\hat{\rho})$ | 0.1797 | 0.1798 | 0.1774 | 0.1771 | 0.1972 | 0.1774 | 0.2213 |
| $bias(\hat{\alpha})$ | -0.0064 | -0.0086 | -0.0066 | -0.0020 | -0.0021 | -0.0027 | 0.0217 |
| $bias(\hat{\beta})$ | 0.0009 | -0.0147 | -0.0006 | -0.0002 | 0.0020 | 0.0021 | 0.0049 |
| $bias(\hat{\sigma}_a^2)$ | -0.0839 | -0.0783 | -0.0820 | -0.0774 | -0.0653 | -0.0897 | 0.1426 |
| $bias(\hat{\sigma}_b^2)$ | -0.0664 | -0.0698 | -0.0730 | -0.0692 | -0.0722 | -0.0835 | -0.0288 |
| $bias(\hat{\rho})$ | 0.0049 | 0.0047 | 0.0047 | 0.0092 | 0.0064 | 0.0089 | 0.0142 |
| $ARE(\hat{\alpha})$ | 0.9879 | 1.1413 | 1.4702 | 1.3697 | 1.0187 | 3.2313 | 1.8647 |
| $ARE(\hat{\beta})$ | 1.7604 | 1.2652 | 1.1873 | 0.7123 | 0.7887 | 0.6345 | 0.8822 |
| $ARE(\hat{\sigma}_a^2)$ | 0.6544 | 0.6561 | 0.6520 | 0.6574 | 0.6224 | 0.6422 | 0.8884 |
| $ARE(\hat{\sigma}_b^2)$ | 0.4686 | 0.4621 | 0.4550 | 0.4549 | 0.4550 | 0.4608 | 0.5045 |
| $ARE(\hat{\rho})$ | 3.4973 | 3.4901 | 3.4663 | 3.4637 | 3.6884 | 3.4618 | 3.9286 |
| Total rank score | 67 | 63 | 51 | 44 | 48 | 50 | 94 |
| Overall rank | 6 | 5 | 4 | 1 | 2 | 3 | 7 |
| Coverage probability | 0.9358 | 0.9334 | 0.9328 | 0.9322 | 0.9374 | 0.9316 | 0.9353 |
| Av. computing time | 8.4471 | 7.3521 | 6.9381 | 9.6094 | 7.4101 | 5.9997 | 58.256 |

**Table 7:** MSE, mean bias, ARE, average computing time in minutes and average total iteration of the estimated values of the five parameters using the coordinate ascent algorithm based on different initial starts at *ρ*=-0.3.

| Criteria | DL | DL2 | HO | HO2 | HM | HS | MLRIV |
| --- | --- | --- | --- | --- | --- | --- | --- |
| $MSE(\hat{\alpha})$ | 2.4732 | 2.5262 | 2.5082 | 2.5166 | 2.5588 | 2.4658 | 2.9165 |
| $MSE(\hat{\beta})$ | 2.4536 | 2.4845 | 2.4795 | 2.4046 | 2.4878 | 2.4108 | 2.6310 |
| $MSE(\hat{\sigma}_a^2)$ | 0.1290 | 0.1205 | 0.1231 | 0.0988 | 0.1174 | 0.1176 | 1.6253 |
| $MSE(\hat{\sigma}_b^2)$ | 0.1310 | 0.1274 | 0.1406 | 0.1153 | 0.1225 | 0.1544 | 0.3092 |
| $MSE(\hat{\rho})$ | 0.1595 | 0.1531 | 0.1535 | 0.1509 | 0.1734 | 0.1546 | 0.2039 |
| $bias(\hat{\alpha})$ | -0.0061 | -0.0071 | -0.0009 | 0.0030 | -0.0001 | -0.0041 | 0.0188 |
| $bias(\hat{\beta})$ | -0.0047 | -0.0057 | -0.0020 | -0.0115 | -0.0032 | -0.0044 | 0.0142 |
| $bias(\hat{\sigma}_a^2)$ | -0.0748 | -0.0839 | -0.0811 | -0.0825 | -0.0599 | -0.0814 | 0.1665 |
| $bias(\hat{\sigma}_b^2)$ | -0.0687 | -0.0730 | -0.0707 | -0.0712 | -0.0727 | -0.0720 | -0.0228 |
| $bias(\hat{\rho})$ | 0.0236 | 0.0221 | 0.0165 | 0.0167 | 0.0293 | 0.0176 | 0.0497 |
| $ARE(\hat{\alpha})$ | 1.5801 | 3.7292 | 0.9986 | 1.1775 | 1.6147 | 1.3351 | 0.8760 |
| $ARE(\hat{\beta})$ | 0.8925 | 3.3295 | 0.9010 | 1.3248 | 0.6368 | 1.2571 | 2.5015 |
| $ARE(\hat{\sigma}_a^2)$ | 0.6528 | 0.6420 | 0.6401 | 0.6370 | 0.6343 | 0.6495 | 0.9358 |
| $ARE(\hat{\sigma}_b^2)$ | 0.4645 | 0.4602 | 0.4572 | 0.4528 | 0.4579 | 0.4696 | 0.5008 |
| $ARE(\hat{\rho})$ | 1.0670 | 1.0447 | 1.0463 | 1.0372 | 1.1189 | 1.0497 | 1.2129 |
| Total rank score | 61 | 71 | 43 | 40 | 55 | 58 | 92 |
| Overall rank | 5 | 6 | 2 | 1 | 3 | 4 | 7 |
| Coverage probability | 0.9415 | 0.9377 | 0.9394 | 0.9389 | 0.9379 | 0.9394 | 0.9365 |
| Av. computing time | 7.1433 | 6.6200 | 8.7898 | 6.1137 | 7.9002 | 7.2197 | 64.2529 |

**Table 8:** MSE, mean bias, ARE, average computing time in minutes and average total iteration of the estimated values of the five parameters using the coordinate ascent algorithm based on different initial starts at *ρ*=-0.5.

| Criteria | DL | DL2 | HO | HO2 | HM | HS | MLRIV |
| --- | --- | --- | --- | --- | --- | --- | --- |
| $MSE(\hat{\alpha})$ | 2.5245 | 2.5555 | 2.5457 | 2.6056 | 4.0167 | 2.5624 | 2.8422 |
| $MSE(\hat{\beta})$ | 2.4202 | 2.4811 | 2.5546 | 2.4806 | 2.4499 | 2.4338 | 2.5231 |
| $MSE(\hat{\sigma}_a^2)$ | 0.1252 | 0.1164 | 0.1141 | 0.1106 | 0.1381 | 0.1135 | 1.4274 |
| $MSE(\hat{\sigma}_b^2)$ | 0.1492 | 0.1312 | 0.1343 | 0.1287 | 0.1228 | 0.1399 | 0.3590 |
| $MSE(\hat{\rho})$ | 0.1159 | 0.1120 | 0.1125 | 0.1199 | 0.1372 | 0.1132 | 0.1725 |
| $bias(\hat{\alpha})$ | -0.0055 | -0.0039 | -0.0096 | -0.0054 | -0.0149 | 0.0011 | 0.0218 |
| $bias(\hat{\beta})$ | -0.0004 | 0.0005 | -0.0079 | -0.0035 | 0.0003 | 0.0017 | 0.0137 |
| $bias(\hat{\sigma}_a^2)$ | -0.0670 | -0.0686 | -0.0742 | -0.0658 | -0.0488 | -0.0645 | 0.1562 |
| $bias(\hat{\sigma}_b^2)$ | -0.0669 | -0.0726 | -0.0786 | -0.0708 | -0.0715 | -0.0711 | -0.0270 |
| $bias(\hat{\rho})$ | 0.0348 | 0.0305 | 0.0384 | 0.0390 | 0.0547 | 0.0296 | 0.0825 |
| $ARE(\hat{\alpha})$ | 1.7965 | 1.1603 | 1.1868 | 0.8828 | 1.2811 | 1.6824 | 1.1621 |
| $ARE(\hat{\beta})$ | 0.6085 | 0.6753 | 0.8003 | 0.8897 | 7.4062 | 0.8663 | 0.6279 |
| $ARE(\hat{\sigma}_a^2)$ | 0.6451 | 0.6420 | 0.6365 | 0.6391 | 0.6352 | 0.6434 | 0.9145 |
| $ARE(\hat{\sigma}_b^2)$ | 0.4663 | 0.4637 | 0.4586 | 0.4536 | 0.4486 | 0.4634 | 0.4875 |
| $ARE(\hat{\rho})$ | 0.5196 | 0.5089 | 0.5110 | 0.5267 | 0.5589 | 0.5172 | 0.6119 |
| Total rank score | 56 | 49 | 61 | 53 | 62 | 51 | 88 |
| Overall rank | 4 | 1 | 5 | 3 | 6 | 2 | 7 |
| Coverage probability | 0.9395 | 0.9397 | 0.9423 | 0.9437 | 0.9458 | 0.9395 | 0.9440 |
| Av. computing time | 10.7919 | 8.2535 | 7.6530 | 8.0867 | 9.2990 | 8.0864 | 53.5529 |

**Table 9:** MSE, mean bias, ARE, average computing time in minutes and average total iteration of the estimated values of the five parameters using the coordinate ascent algorithm based on different initial starts at *ρ*=-0.9.

| Criteria | DL | DL2 | HO | HO2 | HM | HS | MLRIV |
| --- | --- | --- | --- | --- | --- | --- | --- |
| $MSE(\hat{\alpha})$ | 2.5642 | 2.5870 | 2.6619 | 2.5745 | 2.6234 | 2.5551 | 2.8771 |
| $MSE(\hat{\beta})$ | 2.3667 | 2.3820 | 2.5080 | 2.3664 | 2.4653 | 2.4534 | 2.6843 |
| $MSE(\hat{\sigma}_a^2)$ | 0.1916 | 0.1853 | 0.1898 | 0.2268 | 0.1802 | 0.2015 | 0.9187 |
| $MSE(\hat{\sigma}_b^2)$ | 0.2869 | 0.1960 | 0.2458 | 0.2435 | 0.1573 | 0.2785 | 0.4224 |
| $MSE(\hat{\rho})$ | 0.0325 | 0.0345 | 0.0345 | 0.0354 | 0.0419 | 0.0342 | 0.1102 |
| $bias(\hat{\alpha})$ | 0.0125 | 0.0074 | 0.0065 | 0.0003 | 0.0046 | 0.0055 | 0.0220 |
| $bias(\hat{\beta})$ | -0.0012 | -0.0080 | -0.0083 | -0.0034 | 0.0026 | -0.0086 | 0.0169 |
| $bias(\hat{\sigma}_a^2)$ | -0.0118 | -0.0217 | -0.0208 | -0.0004 | -0.0114 | -0.0173 | 0.1548 |
| $bias(\hat{\sigma}_b^2)$ | -0.0514 | -0.0665 | -0.0643 | -0.0567 | -0.0735 | -0.0585 | -0.0269 |
| $bias(\hat{\rho})$ | 0.0557 | 0.0571 | 0.0569 | 0.0568 | 0.0691 | 0.0553 | 0.1629 |
| $ARE(\hat{\alpha})$ | 0.9721 | 0.9000 | 1.0711 | 1.2287 | 1.1388 | 1.3557 | 1.0871 |
| $ARE(\hat{\beta})$ | 0.8938 | 0.7479 | 0.7566 | 0.8744 | 0.9241 | 1.5989 | 0.8269 |
| $ARE(\hat{\sigma}_a^2)$ | 0.6769 | 0.6599 | 0.6737 | 0.6874 | 0.6521 | 0.6692 | 0.8779 |
| $ARE(\hat{\sigma}_b^2)$ | 0.4871 | 0.4770 | 0.4865 | 0.4812 | 0.4597 | 0.4957 | 0.5087 |
| $ARE(\hat{\rho})$ | 0.1064 | 0.1082 | 0.1085 | 0.1070 | 0.1189 | 0.1066 | 0.2100 |
| Total rank score | 47 | 50 | 63 | 51 | 56 | 60 | 92 |
| Overall rank | 1 | 2 | 6 | 3 | 4 | 5 | 7 |
| Coverage probability | 0.9528 | 0.9457 | 0.9465 | 0.9500 | 0.9525 | 0.9476 | 0.9466 |
| Av. computing time | 23.6592 | 18.7869 | 22.8118 | 21.1147 | 19.7893 | 20.7206 | 93.2454 |

**Table 10:** MSE, mean bias, ARE, average computing time in minutes and average total iteration of the estimated values of the five parameters using the coordinate ascent algorithm based on different initial starts for a homogenous population.

| Criteria | DL | DL2 | HO | HO2 | HM | HS | MLRIV |
| --- | --- | --- | --- | --- | --- | --- | --- |
| $MSE(\hat{\alpha})$ | 2.6130 | 2.6683 | 2.5768 | 2.5409 | 2.4671 | 2.5842 | 2.8403 |
| $MSE(\hat{\beta})$ | 2.5169 | 2.5354 | 2.4285 | 2.4533 | 2.3385 | 2.4506 | 2.5863 |
| $MSE(\hat{\sigma}_a^2)$ | 0.0954 | 0.0519 | 0.0789 | 0.0482 | 0.0411 | 0.0441 | 1.5517 |
| $MSE(\hat{\sigma}_b^2)$ | 0.0360 | 0.0475 | 0.0240 | 0.0290 | 0.0326 | 0.0419 | 0.6482 |
| $MSE(\hat{\rho})$ | 0.0980 | 0.1004 | 0.1004 | 0.1009 | 0.1000 | 0.1028 | 0.1878 |
| $bias(\hat{\alpha})$ | 0.0004 | 0.0012 | 0.0109 | 0.0050 | 0.0040 | 0.0048 | 0.0221 |
| $bias(\hat{\beta})$ | 0.0110 | 0.0089 | 0.0085 | 0.0117 | 0.0074 | 0.0090 | 0.0250 |
| $bias(\hat{\sigma}_a^2)$ | 0.0133 | 0.0099 | 0.0146 | 0.0090 | 0.0057 | 0.0105 | 0.2591 |
| $bias(\hat{\sigma}_b^2)$ | 0.0116 | 0.0115 | 0.0080 | 0.0095 | 0.0044 | 0.0108 | 0.0950 |
| $bias(\hat{\rho})$ | 0.0451 | 0.0433 | 0.0434 | 0.0422 | 0.0466 | 0.0457 | 0.1166 |
| $ARE(\hat{\alpha})$ | 0.7707 | 0.7426 | 0.6855 | 0.6688 | 0.6455 | 0.9661 | 1.1121 |
| $ARE(\hat{\beta})$ | 1.1527 | 0.3003 | 0.2921 | 0.3281 | 0.3730 | 0.3402 | 5.8159 |
| $ARE(\hat{\sigma}_a^2)$ | 1.0979 | 1.0619 | 1.1131 | 1.0592 | 1.0538 | 1.0590 | 2.7876 |
| $ARE(\hat{\sigma}_b^2)$ | 1.0190 | 1.0237 | 0.9448 | 0.9673 | 0.9455 | 0.9982 | 2.2563 |
| $ARE(\hat{\rho})$ | 0.4732 | 0.4794 | 0.4782 | 0.4781 | 0.4734 | 0.4805 | 0.6245 |
| Total rank score | 64 | 61 | 48 | 47 | 31 | 63 | 105 |
| Overall rank | 6 | 4 | 3 | 2 | 1 | 5 | 7 |
| Coverage probability | 0.9725 | 0.9707 | 0.9716 | 0.9716 | 0.9735 | 0.9714 | 0.9762 |
| Av. computing time | 9.1498 | 6.6450 | 7.5045 | 6.7165 | 8.0233 | 7.0037 | 53.1311 |

**Table 11:**
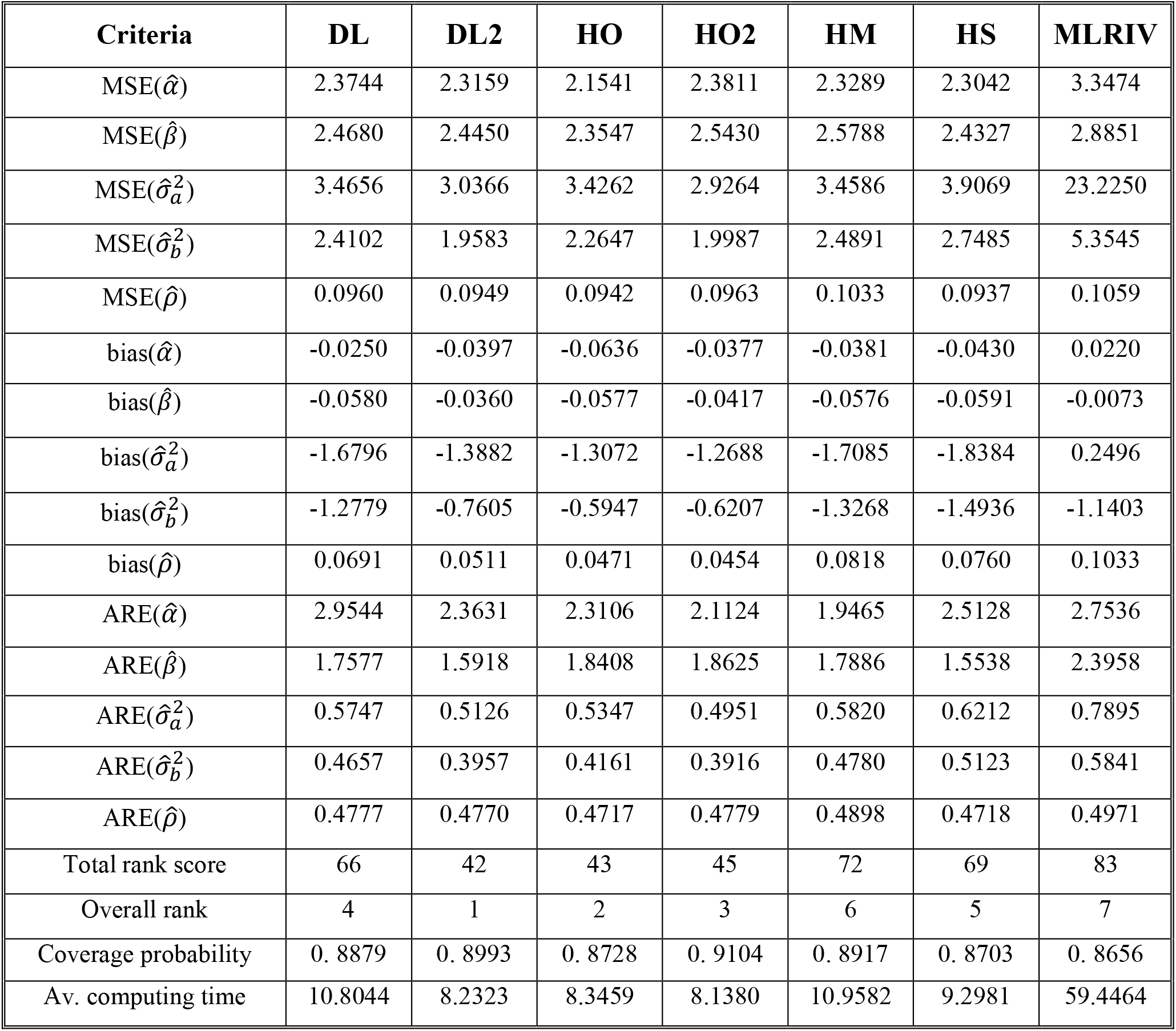
MSE, mean bias, ARE, average computing time in minutes and average total iteration of the estimated values of the five parameters using the coordinate ascent algorithm based on different initial starts for heterogeneous population.

From the tables it shows that there are few advantages in using the MLRIV over closed form methods to provide initial values to the co-ordinate ascent algorithm. Although in one instance the MLRIV estimator for 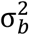 had the least absolute bias in 9 of the 11 scenarios, overall it performed poorly compared with the closed form methods. In all the scenarios investigated the MLRIV estimators for the 5 parameters had the lowest overall rank (highest score) in terms of MSE, absolute bias and ARE by a substantial margin.

The MSE for the MLRIV estimates for the 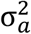 and 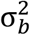 is also of note. This is consistently larger (sometimes by a factor of 10) than the corresponding MSE for the estimates from the closed form methods. The box and whisker plots (see appendix) gives a clearer picture of the distribution of estimates for the 5 parameters. Here it can be seen that there is substantially wider variation in the MLRIV estimates for 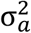 and 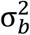 across all scenarios than those provided by the closed form estimators.

Furthermore the computation time is substantially longer when using the MLRIV. For example for *k*=10 studies the average computation time using the closed form methods is between 11½ and 13½ minutes compared with 66 minutes for the MLRIV. Thus, using the closed form methods to provide initial values to the algorithm reduces the average computation time by at least 80% compared with the MLRIV.

Comparing the closed form methods in terms of MSE, bias and ARE no one initial estimator dominates the others over all 5 parameters. For individual parameters, the HM estimator for 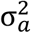 had the lowest absolute bias and ARE in 7 and 8 of the 11 scenarios, respectively. However, in general, the highest rank was more evenly distributed across the closed form methods.

The coverage probabilities of the confidence ellipses for *α* and *β* are estimated using methods described in [25] and have been previously used in [8]. The 95% coverage probability was mostly above 93% for all methods across the different scenarios but for *k*=5 and the ‘homogeneous’ population, (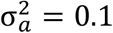 and 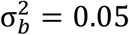), the coverage was above 97%. In contrast, for the ‘heterogeneous’ population (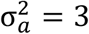and 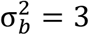) coverage was as low as 87% for some of the closed form methods and highest for the HO2 estimator where it was 91%.

In table 12, the distributions of the scores for different methods are summarised. Although no closed form was clearly better than the others, HO and HO2 have lowest median and second lowest minimum scores. The upper quartiles for HO2 and HO are 53 and 56.5 and these are the lowest and second lowest upper quartile. The next closest is HS with 59. The mean (standard deviation) of the total rank score for HO2 is 50.5 (6.6). This is the lowest mean and second lowest standard deviation amongst the closed form methods.

**Table 12:** Summary of distributions of total rank scores

|  | DL | DL2 | HO | HO2 | HM | HS | MLRIV |
| --- | --- | --- | --- | --- | --- | --- | --- |
| Minimum | 47 | 42 | 40 | 40 | 31 | 49 | 83 |
| Lower quartile | 50 | 49.5 | 45.5 | 46 | 53.5 | 50.5 | 92 |
| Median | 56 | 56 | 51 | 51 | 56 | 55 | 93 |
| Upper Quartile | 62.5 | 61.5 | 56.5 | 53 | 60.5 | 59 | 94 |
| Maximum | 67 | 71 | 63 | 65 | 72 | 69 | 105 |
| Mean | 56.2 | 55.5 | 51.4 | 50.5 | 55.7 | 55.6 | 92.9 |
| Standard deviation | 7.3 | 8.5 | 7.5 | 6.6 | 10.4 | 6.4 | 5.3 |

## 8. Discussion

Previous research has demonstrated the effectiveness of using a Newton-Raphson based algorithm to fit the bivariate random effects model for test accuracy meta-analyses [8]. Specifically, when optimised to the BRM, probability of convergence near 100% may be achieved without loss of accuracy of the parameter estimates. An important element of Newton-Raphson based algorithms, as formalised by Kantorovich’s theorem, is the locating of starting points close enough to the global maximum so that the function is well behaved in terms of continuity and boundedness [10]. Although it is difficult to translate this theorem directly into an iterative algorithm it does reinforce the importance of having reasonable initial estimates.

To this end, robust initial values based on maximum likelihood were proposed as starting values [8]. However, they are computationally expensive. Furthermore, it was not clear whether they produced estimates which were any less biased or had lower mean square errors than those obtained from using starting values derived from closed form methods.

In this paper we explored six closed form methods for producing initial estimators to the Newton-Raphson algorithm for maximising the likelihood of the BRM. The immediate benefit of using such methods is that the average computation time for fitting the model for 10 included studies was reduced by around 80% from 66 minutes to between 11½ and 13½ minutes. Of course as the number of included studies increases the computation time increases leading to times in excess of 1 hour even when closed form initial estimators are used. This is a consequence of using the interpretative language, R to run the algorithms which benefits from being illustrative but is slow in comparison to low-level compiled languages such as C. Any further improvements in the computation time of the algorithm used here are likely to require its translation into C/C++.

To summarise the data for each of the estimators the estimates for each of parameters were ranked in terms of the mean squared error (MSE), absolute bias and average relative error (ARE) and then summed. Thus, the highest ranked estimator for a particular scenario had the lowest total rank score. From this it was clear that the closed form methods, in general, performed substantially better than the MLRIV. Indeed the worst closed form methods had a median total rank score that was 37 points lower than the MLRIV.

Although the simulation study demonstrated that no one closed form method dominated the others across all metrics, the distribution of total rank scores suggests that overall the two-step Hedges and Olkin (HO2) estimator may have a modest advantage in terms of accuracy and consistency. However, more extensive analyses over a greater range of scenarios are required before it may be concluded whether this observation is real or not.

Future studies may evaluate other non-Newton-Raphson based optimisation algorithms for the bivariate random effects model used in meta-analysis of test accuracy studies to see if they may be fine-tuned to improve on the performance characteristics whilst matching the probability of convergence of the Newton-Raphson algorithm used here. Furthermore the validity of the BRM in providing estimates that have clinical utility should be subject to investigation [26,27].

In summary, we have demonstrated that the use of closed form methods to estimate the initial values of the Newton-Raphson algorithm to fit the bivariate random effects model substantially improves computational efficiency and reduces the time to convergence. Furthermore it is clear that the closed form methods, in general, improve upon the MLRIV method in other performance characteristics. Overall, the two-step Hedges-Olkin estimator ranked highest in terms of median and mean total rank scores but further research is required to ascertain the significance of this result.

## Acknowledgments

The computations described in this paper were performed using the University of Birmingham’s BlueBEAR HPC service and CaStLeS (Compute and Storage for Life Sciences) resources [28] that provide a High Performance Computing service to the University’s research community. See http://www.birmingham.ac.uk/bear for more details.

