## Supplementary figures for "Optimising a coordinate ascent algorithm for the meta-analysis of test accuracy studies"

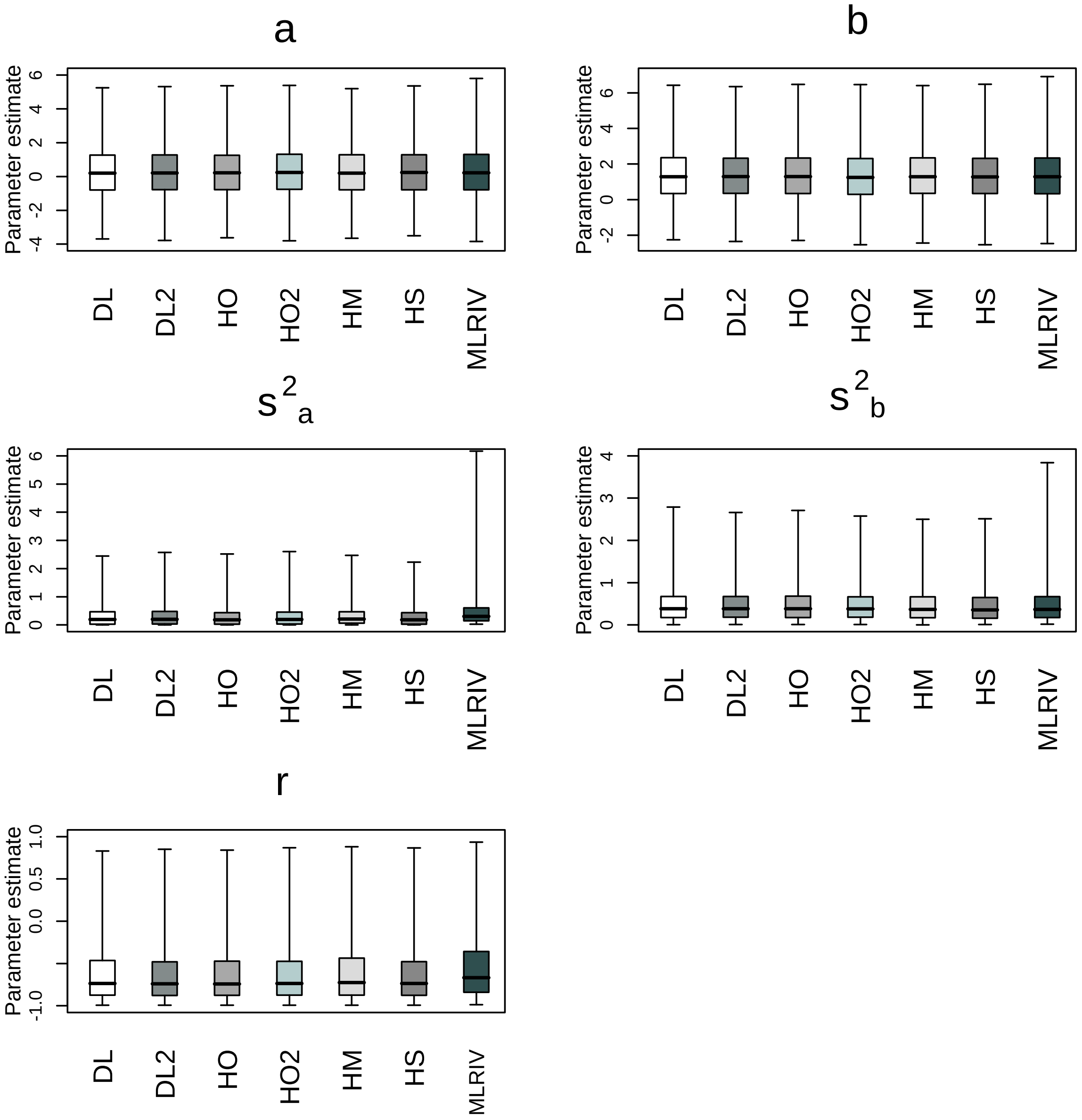


Figure 1: Boxplots of the estimated values of the five parameters obtained from 10000 simulated data using 7 methods at k=5 based on $\sigma_a^2 = 0.4$; $\sigma_b^2 = 0.6$; and $\rho = -0.7$. The boxes are bounded by the lower and upper quartiles and the caps to the whiskers represent the 0.5 and 99.5 percentiles.

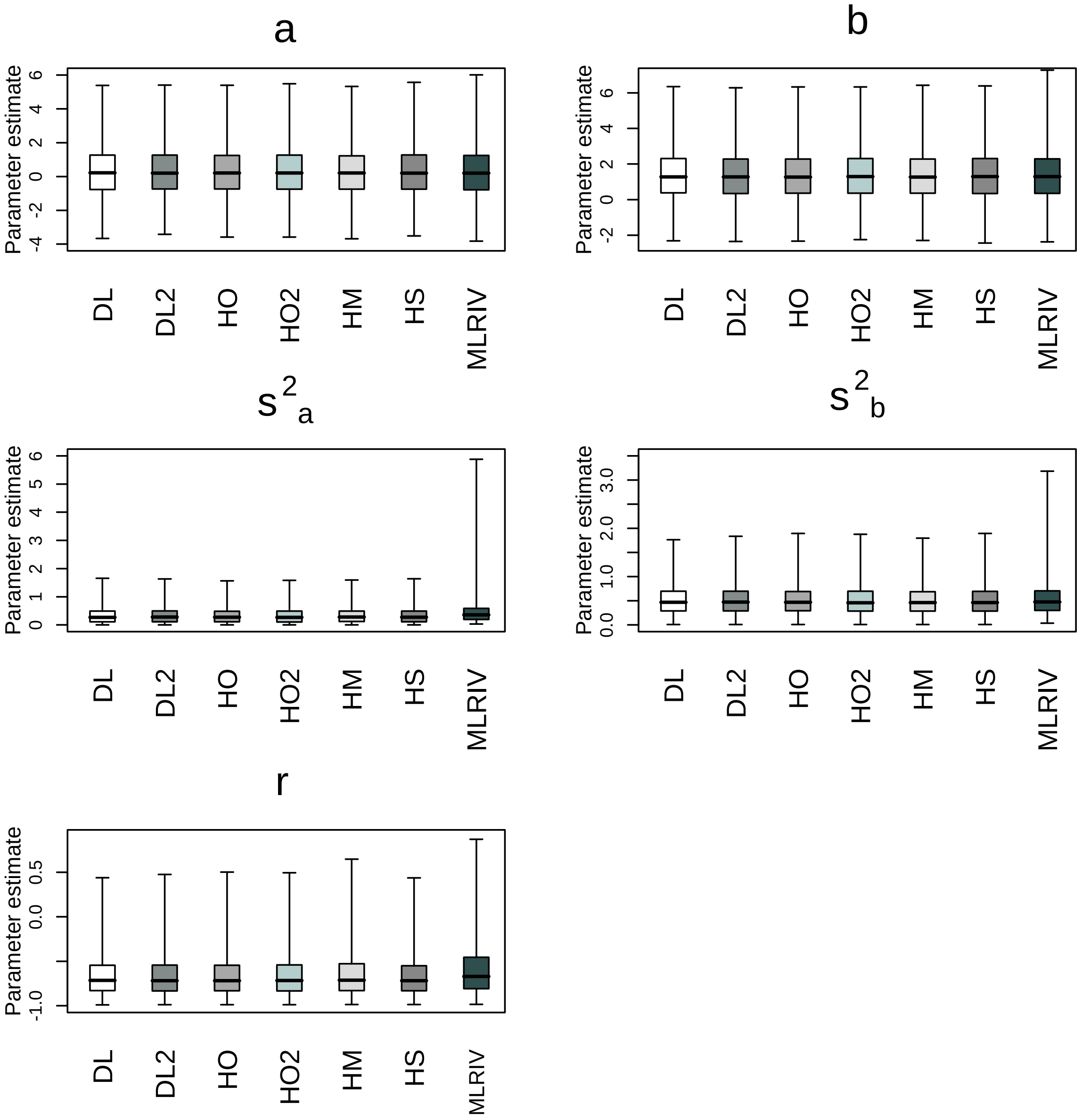


Figure 2: Boxplots of the estimated values of the five parameters obtained from 10000 simulated data using 7 methods at k=10 based on $\sigma_a^2 = 0.4$; $\sigma_b^2 = 0.6$; and $\rho = -0.7$. The boxes are bounded by the lower and upper quartiles and the caps to the whiskers represent the 0.5 and 99.5 percentiles.

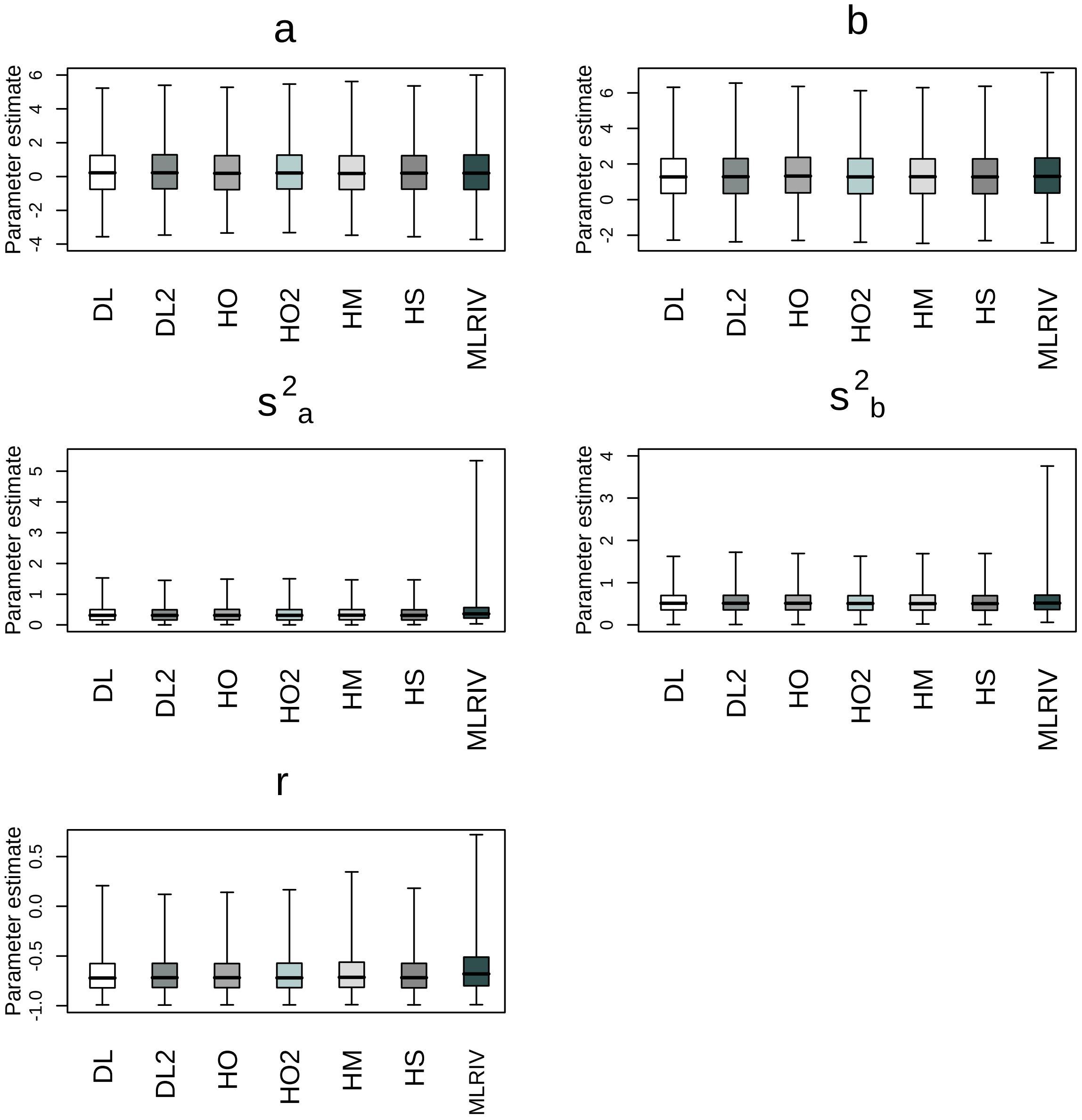


Figure 3: Boxplots of the estimated values of the five parameters obtained from 10000 simulated data using 7 methods at k=15 based on $\sigma_a^2 = 0.4$; $\sigma_b^2 = 0.6$; and $\rho = -0.7$. The boxes are bounded by the lower and upper quartiles and the caps to the whiskers represent the 0.5 and 99.5 percentiles.

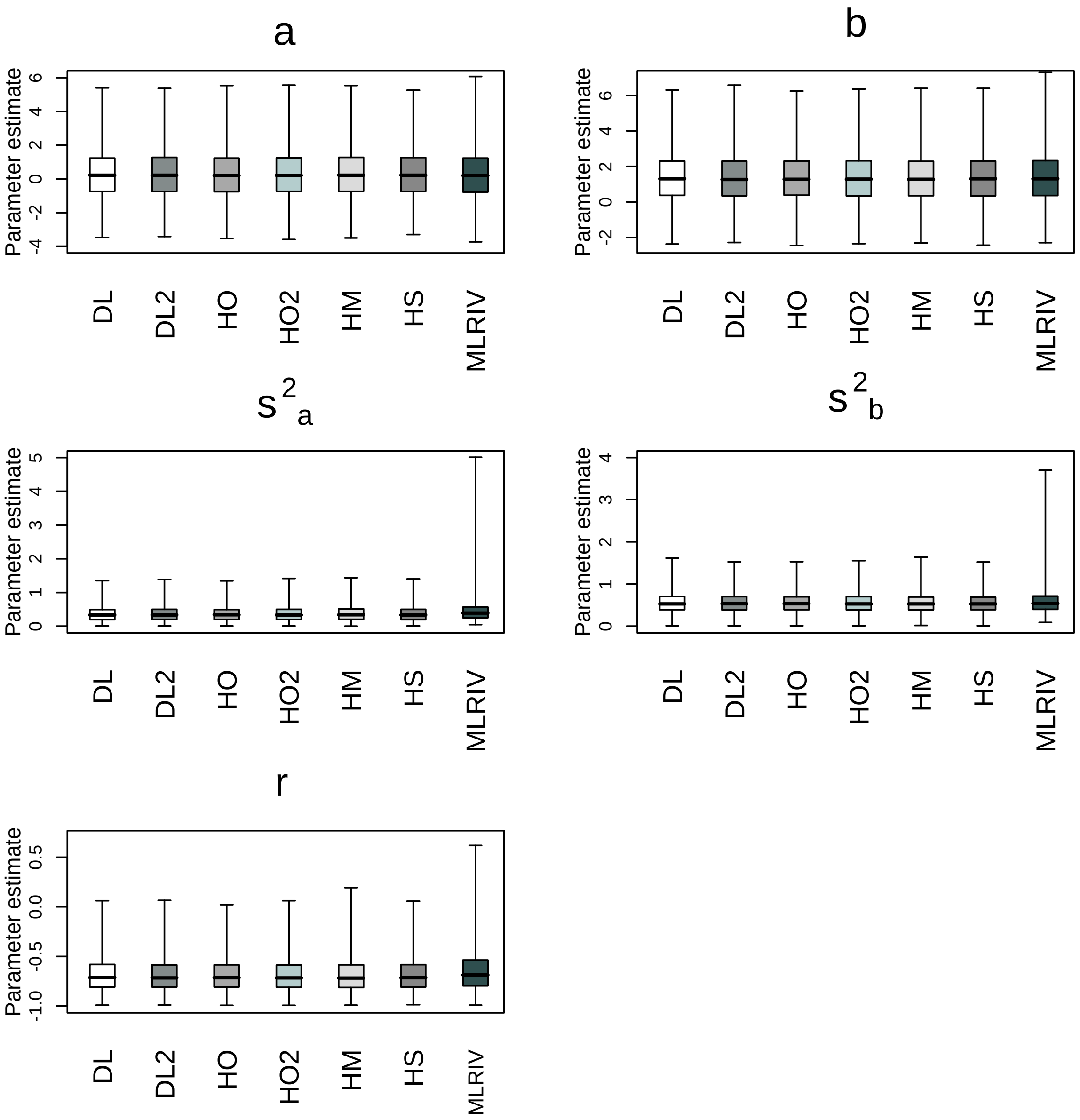


Figure 4: Boxplots of the estimated values of the five parameters obtained from 10000 simulated data using 7 methods at k=20 based on $\sigma_a^2 = 0.4$; $\sigma_b^2 = 0.6$; and $\rho = -0.7$. The boxes are bounded by the lower and upper quartiles and the caps to the whiskers represent the 0.5 and 99.5 percentiles.

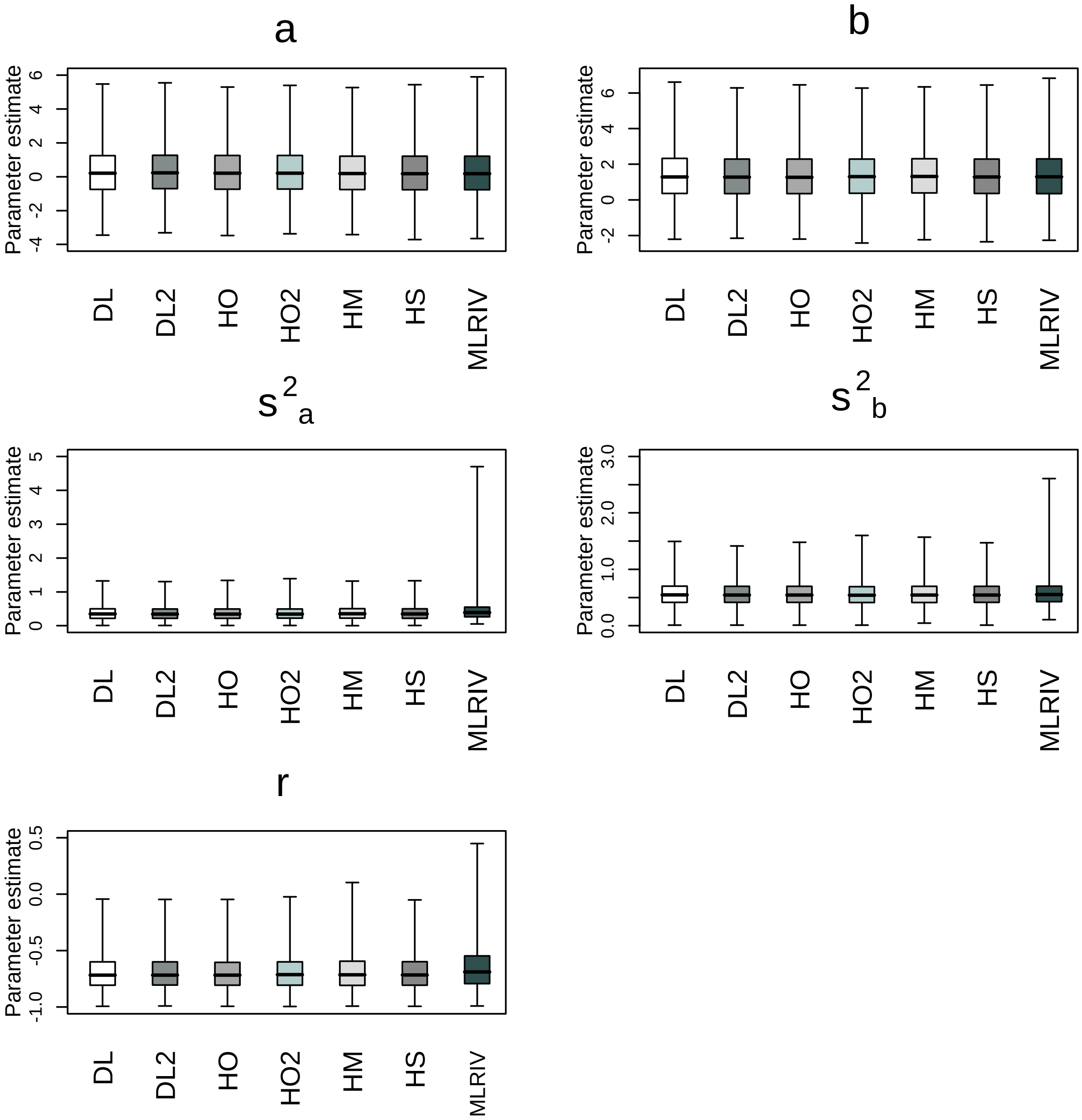


Figure 5: Boxplots of the estimated values of the five parameters obtained from 10000 simulated data using 7 methods at k=25 based on $\sigma_a^2 = 0.4$; $\sigma_b^2 = 0.6$; and $\rho = -0.7$. The boxes are bounded by the lower and upper quartiles and the caps to the whiskers represent the 0.5 and 99.5 percentiles.

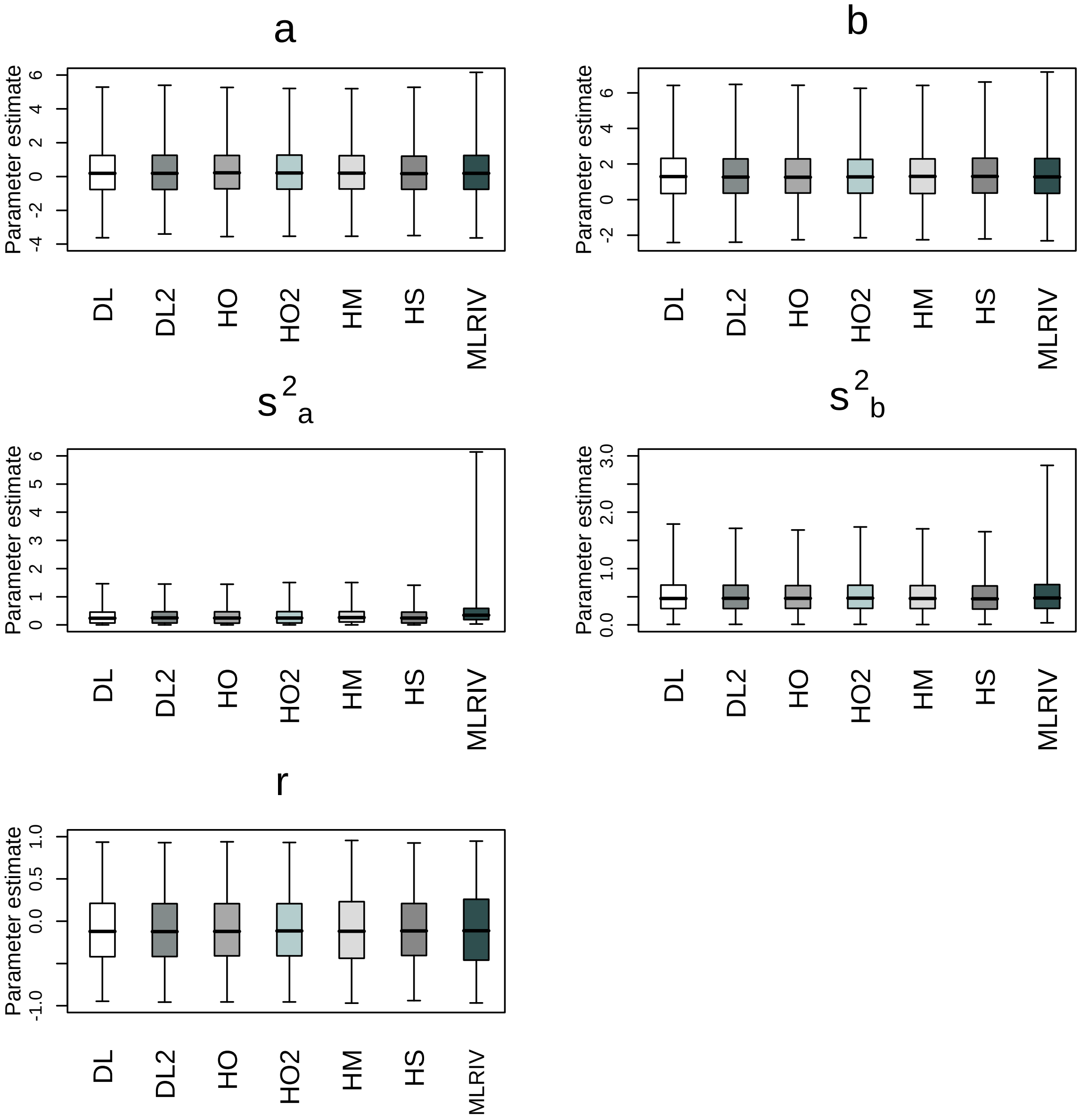


Figure 6: Boxplots of the estimated values of the five parameters obtained from 10000 simulated data using 7 methods at k=10 based on $\sigma_a^2 = 0.4$; $\sigma_b^2 = 0.6$; and $\rho = -0.1$. The boxes are bounded by the lower and upper quartiles and the caps to the whiskers represent the 0.5 and 99.5 percentiles.

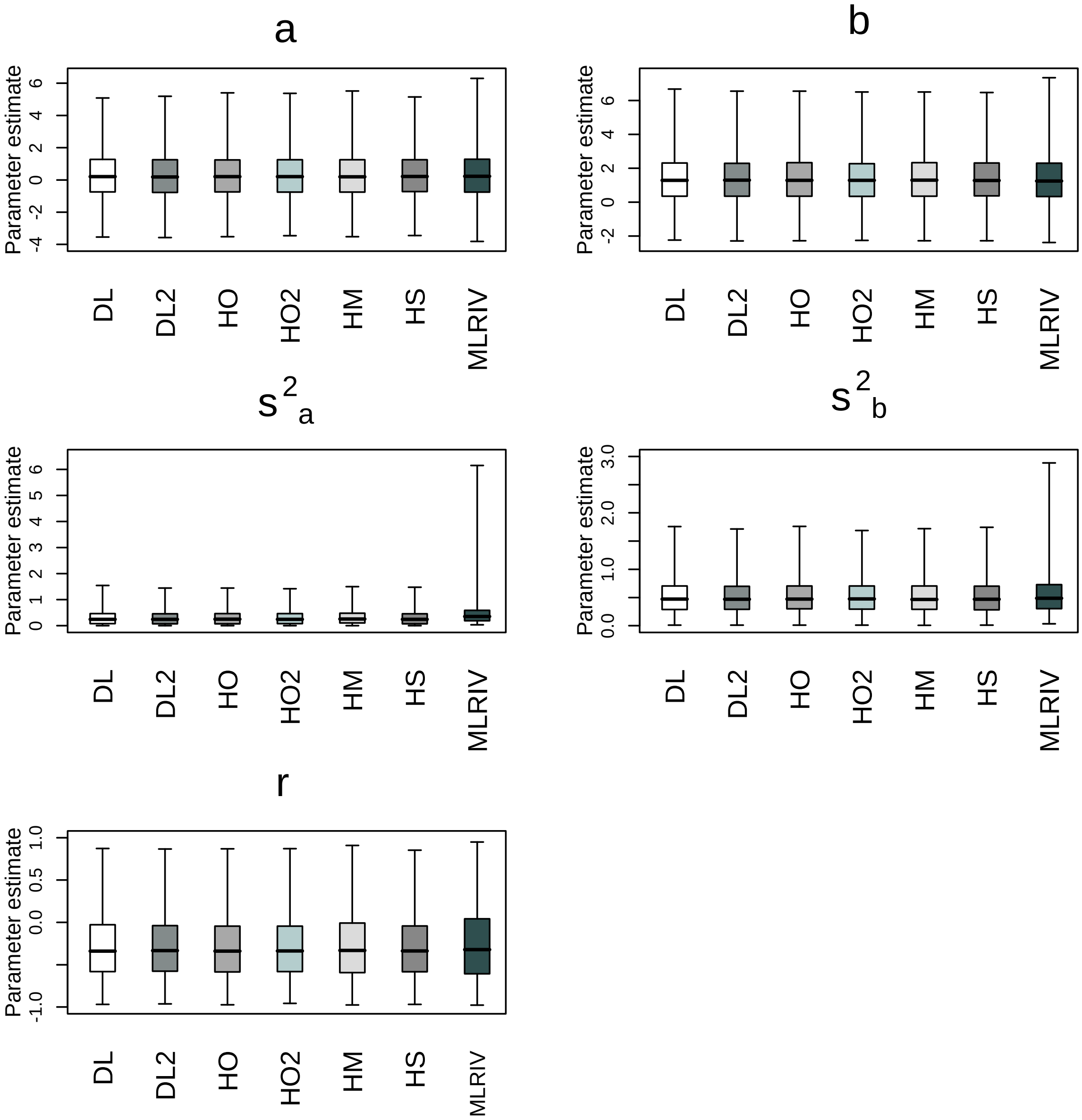


Figure 7: Boxplots of the estimated values of the five parameters obtained from 10000 simulated data using 7 methods at k=10 based on $\sigma_a^2 = 0.4$; $\sigma_b^2 = 0.6$; and $\rho = -0.3$. The boxes are bounded by the lower and upper quartiles and the caps to the whiskers represent the 0.5 and 99.5 percentiles.

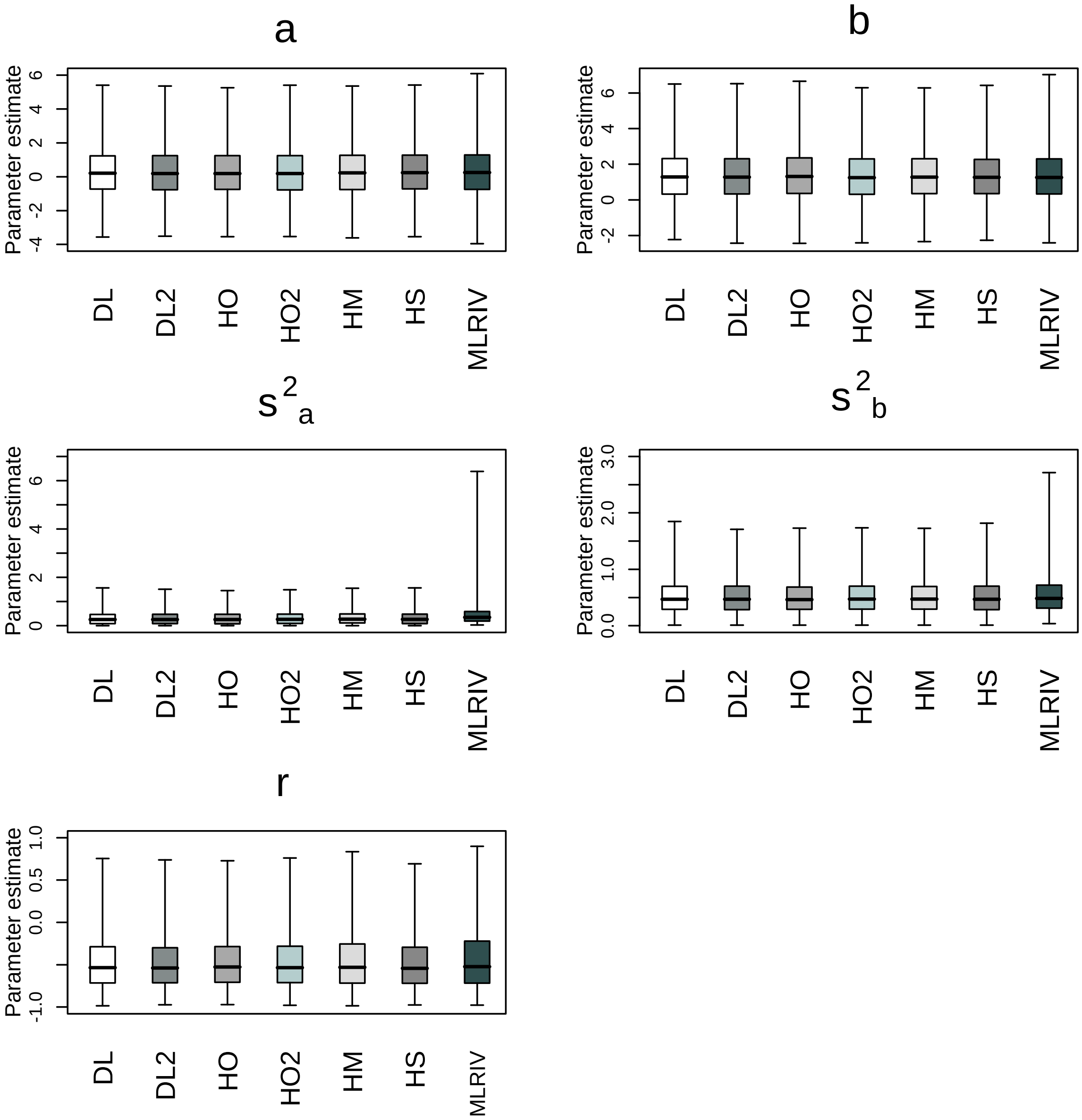


Figure 8: Boxplots of the estimated values of the five parameters obtained from 10000 simulated data using 7 methods at k=10 based on $\sigma_a^2 = 0.4$; $\sigma_b^2 = 0.6$; and $\rho = -0.5$. The boxes are bounded by the lower and upper quartiles and the caps to the whiskers represent the 0.5 and 99.5 percentiles.

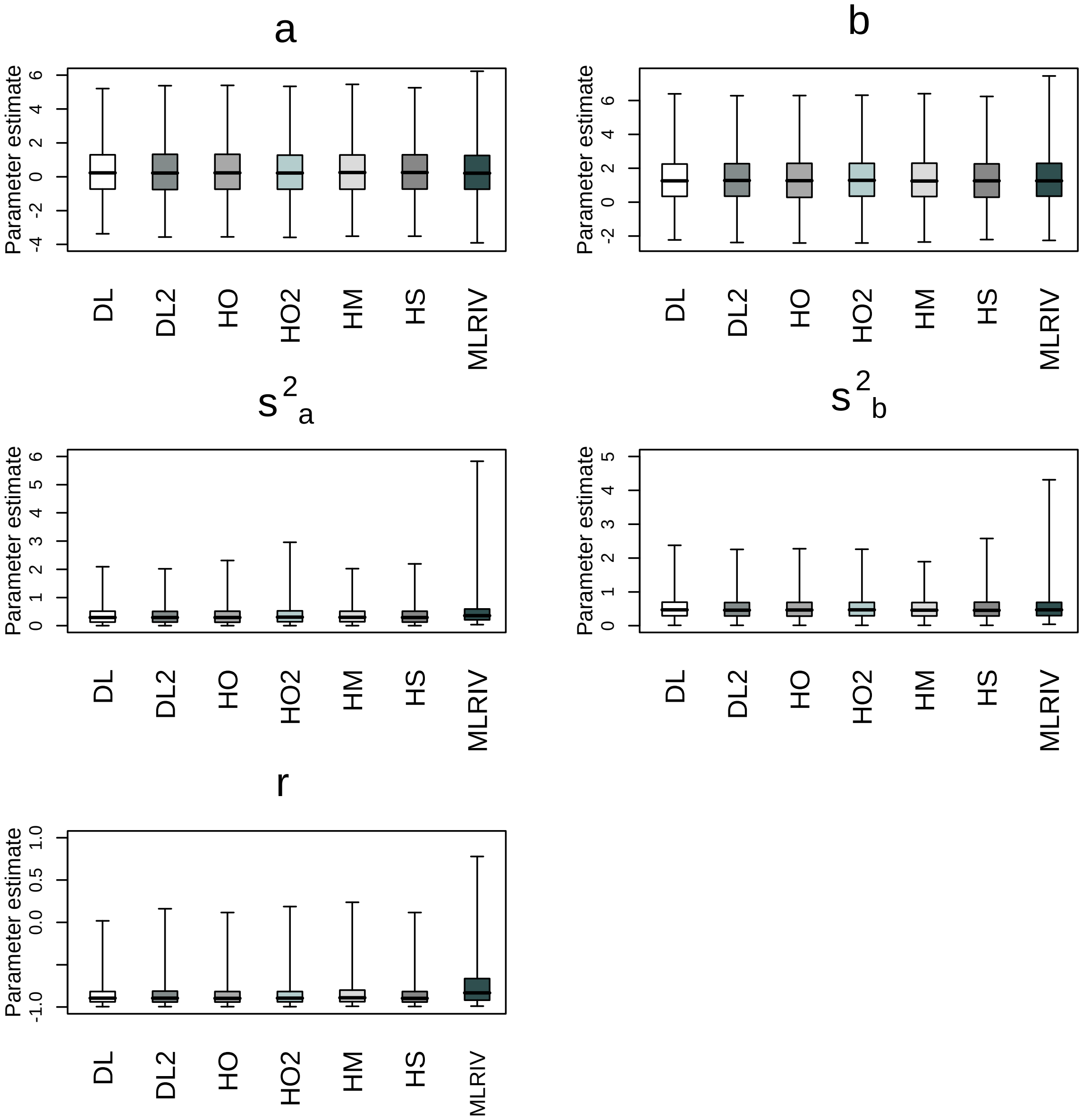


Figure 9: Boxplots of the estimated values of the five parameters obtained from 10000 simulated data using 7 methods at k=10 based on $\sigma_a^2 = 0.4$; $\sigma_b^2 = 0.6$; and $\rho = -0.9$. The boxes are bounded by the lower and upper quartiles and the caps to the whiskers represent the 0.5 and 99.5 percentiles.

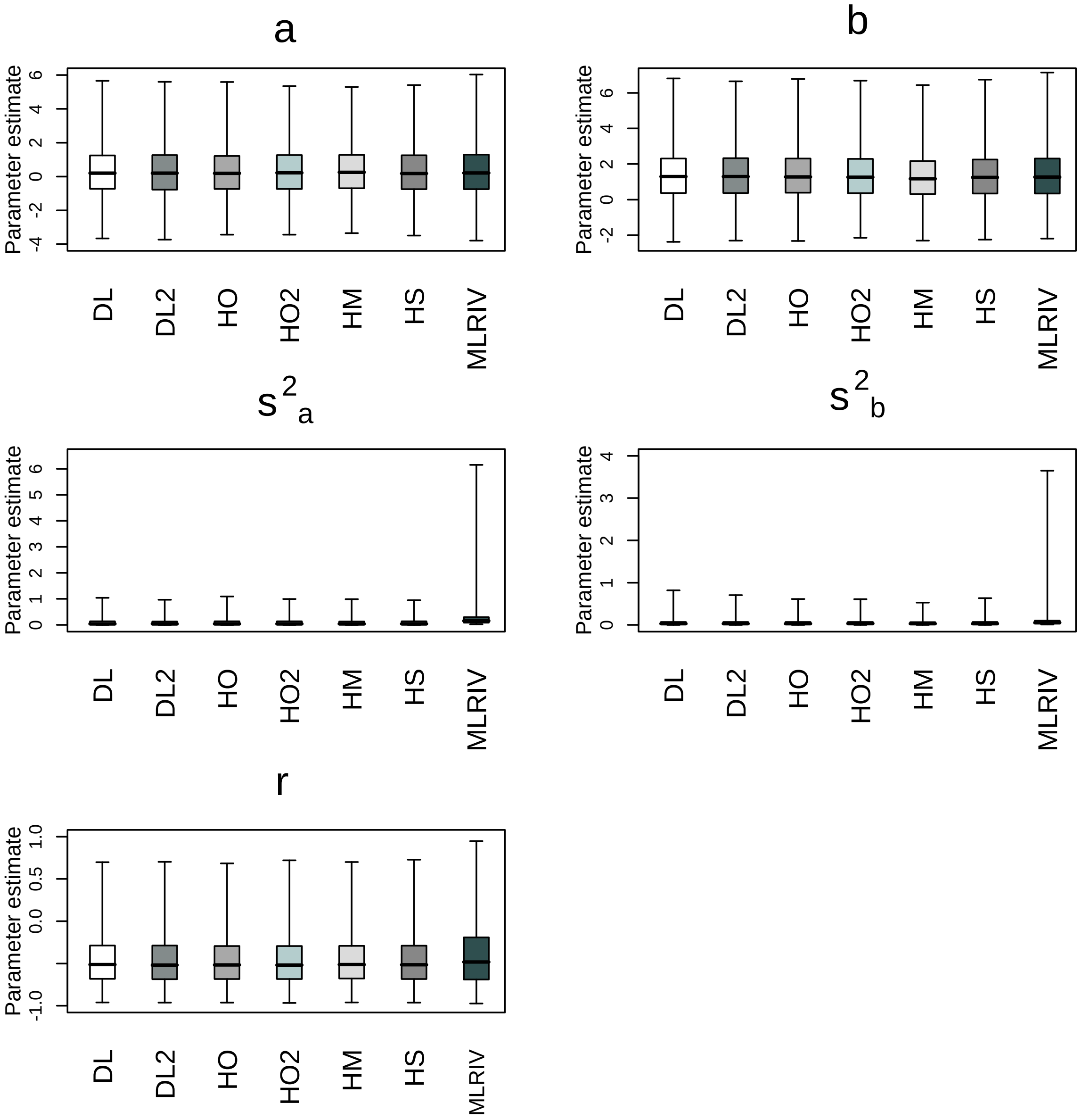


Figure 10: Boxplots of the estimated values of the five parameters obtained from 10000 simulated data using 7 methods at k=10 based on undispersed variances $\sigma_a^2 = 0.1$; $\sigma_b^2 = 0.05$; and $\rho = -0.5$. The boxes are bounded by the lower and upper quartiles and the caps to the whiskers represent the 0.5 and 99.5 percentiles.

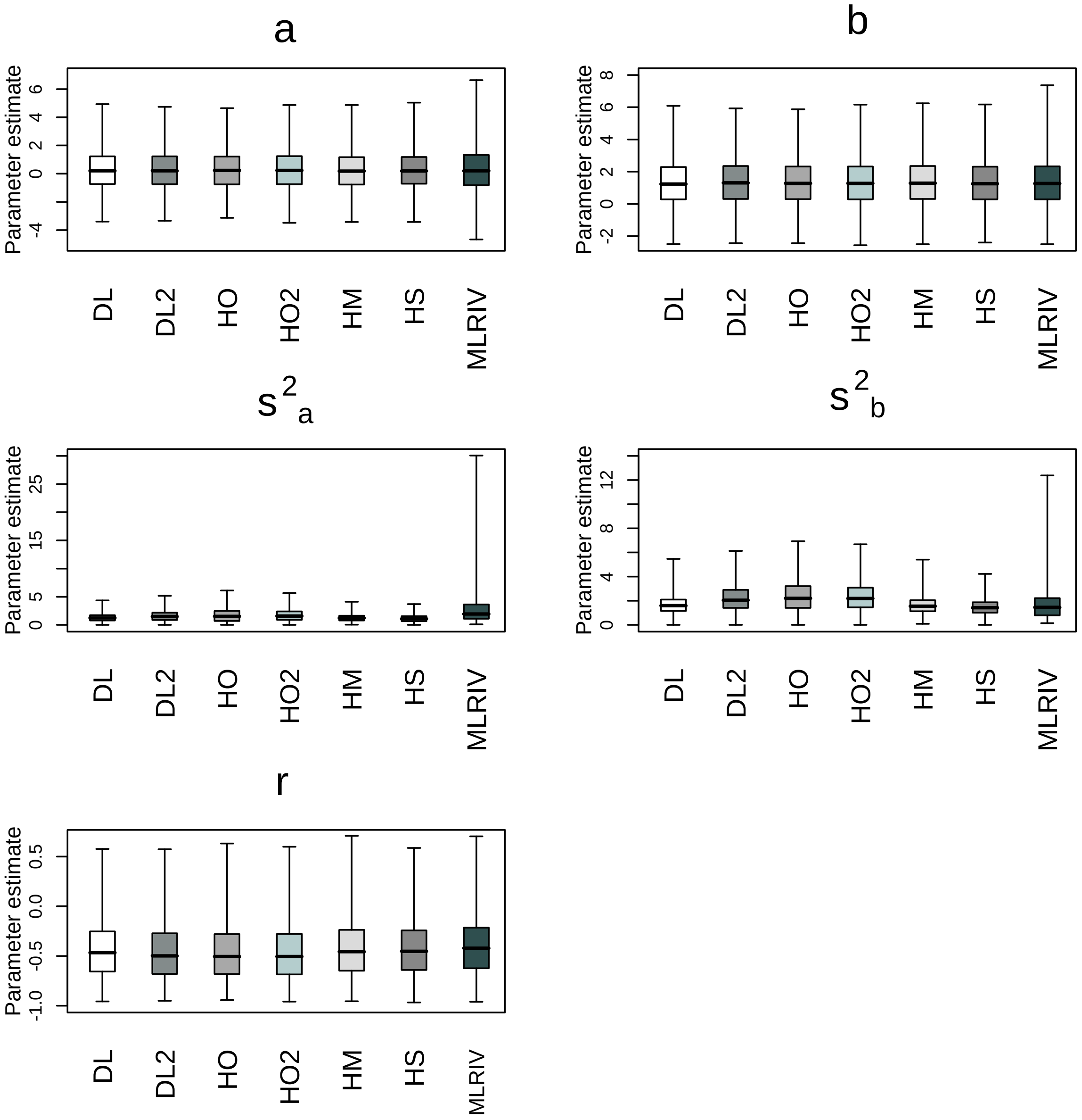


Figure 11: Boxplots of the estimated values of the five parameters obtained from 10000 simulated data using 7 methods at k=10 based on dispersed variances $\sigma_a^2 = 3$; $\sigma_b^2 = 3$; and $\rho = -0.5$. The boxes are bounded by the lower and upper quartiles and the caps to the whiskers represent the 0.5 and 99.5 percentiles.
